# Arginine-Vasopressin Expressing Neurons in the Murine Suprachiasmatic Nucleus Exhibit a Circadian Rhythm in Network Coherence *In Vivo*

**DOI:** 10.1101/2021.12.07.471437

**Authors:** Adam Stowie, Zhimei Qiao, Daniella Do Carmo Buonfiglio, Delaney M. Beckner, J. Christopher Ehlen, Morris Benveniste, Alec J. Davidson

## Abstract

The Suprachiasmatic Nucleus (SCN) is composed of functionally distinct sub-populations of GABAergic neurons which form a neural network responsible for synchronizing most physiological and behavioral circadian rhythms in mammals. To date, little is known regarding which aspects of SCN rhythmicity are generated by individual SCN neurons, and which aspects result from neuronal interaction within a network. Here, we utilize *in vivo* miniaturized microscopy to measure fluorescent GCaMP-reported calcium dynamics in AVP-expressing neurons in the intact SCN of awake, behaving mice. We report that SCN AVP neurons exhibit periodic, slow calcium waves which we demonstrate, using *in vivo* electrical recordings, likely reflect burst-firing. Further, we observe substantial heterogeneity of function in that AVP neurons exhibit unstable rhythms, and relatively weak rhythmicity at the population level. Network analysis reveals that correlated cellular behavior, or coherence, among neuron pairs also exhibited stochastic rhythms with about 33% of pairs rhythmic at any time. Unlike single-cell variables, coherence exhibited a strong rhythm at the population level with time of maximal coherence among AVP neuronal pairs at CT/ZT 6 and 9, coinciding with the timing of maximal neuronal activity for the SCN as a whole. These results demonstrate robust circadian variation in the coordination between stochastically rhythmic neurons and that interactions between AVP neurons in the SCN may be more influential than single-cell activity in the regulation of circadian rhythms. Furthermore, they demonstrate that cells in this circuit, like those in many other circuits imaged *in vivo*, exhibit profound heterogenicity of function over time and space.

**Significance Statement:** This work is the first to employ two novel *in vivo* recording techniques, miniaturized calcium microscopy and optogentically-targeted single unit activity recording, to examine the rhythmic behavior of AVP expressing neurons both at the individual neuronal and network level. These results suggest that while AVP neurons are important for organismal rhythmicity, individual cellular rhythms are unstable and diverse. However, we observed correlated activity among these neurons which appears more reliably rhythmic, suggesting that emergent network properties of the SCN may be more relevant for organismal rhythmicity than individual neuronal characteristics.

## Introduction

In mammals, circadian rhythms are synchronized by the suprachiasmatic nucleus (SCN) of the ventral hypothalamus[1, 2]. At the molecular level, circadian rhythms are driven by a transcription-translation feedback loop in which the core clock genes CLOCK and BMAL1 promote the transcription of the CRY and PER genes, which in turn inhibit their own transcription in a process that takes approximately 24 hours[3]. While most mammalian cells contain this molecular machinery[4], with a few notable exceptions[5, 6], rhythmicity is lost after only a few cycles when isolated from the SCN[7]. SCN neurons, cultured in isolation or experimentally separated, exhibit weak or unstable rhythms[4, 8–10]. Thus, although molecular clock machinery is necessary for the generation of circadian rhythms, individual neurons are insufficient to maintain longitudinal rhythms. There is increasing evidence that the neurons in the SCN act as a neural network in which interactions between neurons within the circuit may be responsible for regulation circadian rhythms. SCN brain slices exhibit robust rhythms in core clock gene expression and intracellular calcium dynamics *ex vivo* for weeks to months[7, 11], and these rhythms are reversibly dampened when cellular communication is disrupted with TTX[11–13]. In addition, longitudinal bioluminescent slice recordings reveal the existence of an organized wave of clock gene expression which propagates through the SCN in a stereotypic fashion, for both *mPer1*[14] and *mPer2*[11, 15]. This temporal organization is modified in a predictable and reversible way by alterations in daylength, suggesting that this emergent property of the network encodes important information about the external world[16, 17], and that rhythmic function among this population is typified by heterogeneity.

Most SCN neurons are GABAergic[18], but can be sub-divided based on other expressed proteins. Distinct populations of vasoactive intestinal polypeptide (VIP)-expressing, gastrin releasing peptide (GRP)-expressing and arginine vasopressin (AVP)-expressing GABAergic neurons are somewhat divided anatomically [19]. In addition, approximately 40% of SCN neurons express the neuropeptide Neuromedin S (NMS) and approximately 60% express the dopamine receptor DRD1[20]; however, these genes are not expressed in distinct neuronal populations and these neurons may also express AVP or VIP[21].

Activity of these neuronal subpopulations may correspond to different roles within the circadian circuit. For instance, loss of NMS neurons abolishes rhythmicity entirely[21]. VIP- and GRP-expressing neurons likely receive input from the retina and play a role in entrainment of the SCN to the light/dark cycle[22, 23]. Also, VIP and GABA act together to maintain synchrony between SCN neurons, particularly under changing environmental lighting[16]. However, the role of AVP neurons is less clear. Disrupting the circadian clock by selectively knocking out either *bmal1* or *casein kinase 1 delta* in AVP neurons lengthened the period of oscillation[24–26], although selective destruction of AVP neurons themselves does not affect locomotor activity rhythms[27]. Taken together, available data suggest that rhythmicity of the SCN network is rather resistant to the loss of specific neuronal populations of the SCN network.

Multi-unit neuronal activity (MUA) recordings show that the intact SCN network exhibits a robust firing rhythm peaking during the day, even in constant darkness [28–31]. Cell-type specific contributions to circadian timing in the intact SCN is less understood. Measurement of VIP-neuron gene expression and intracellular calcium via fiber photometry (FP) show robust population level rhythms in calcium as well as Per1, Per2, and Cry1 expression *in vivo*[23, 32]. This approach highlights the importance of VIP neurons for photic entrainment[23, 33], and the importance of GABA release from AVP neurons in regulating the timing of firing of other SCN neurons[34]. However, such population measures do not provide the cellular resolution needed to elucidate how individual neurons interact in neural networks to exact function and behavior. Without the measure of individual neuron activity, we cannot know how diverse neuronal behavior may be within a network.

Studies on neural networks have indicated that cohesive function of the circuit can occur with significant heterogeneity of function among individual neurons[35–38]. New *in vivo* fluorescent approaches have been developed in which individual neurons of specific cell types within a network can be targeted and recorded within an intact circuit of an awake, behaving animal. Recording from brain slices which lack inputs and outputs, or recording population-level behavior, even *in vivo* (e.g., MUA or FP), could mask potential heterogeneity which may be important for circuit function. State changes in such circuits are a point at which heterogeneity of function can switch to more homogenous, coordinated function. One example is cortical reorganization during sleep, where diverse signaling in wake changes to more coordinated activity during sleep[39]. The mammalian SCN undergoes reliable, repeating state changes each day making it an excellent model for determining how individual neurons contribute to a cohesive behavioral outcome. In slice recordings, significant heterogeneity in the phase of individual cells is observed [40] and this heterogeneity *in vivo* may influence behavior-level functional outcomes [41]. Different cell types exhibit diverse characteristics [26, 42] and a single cell can exhibit stochastic rhythmicity [43], another type of heterogeneity observed within individual cells over time. However, whether such diversity of function across cells and time exists *in vivo*, in an unperturbed SCN circuit is unknown.

In the present work, we characterize bursting electrical behavior of AVP neurons within the intact SCN network using optogenetically-targeted single-unit activity recording (SUA^AVP^). Further, we employ miniaturized fluorescence microscopy to characterize a portion of the SCN network by conducting longitudinal recording of intracellular calcium dynamics in AVP-expressing neurons in the fully intact SCN of freely behaving mice. We report that individual AVP neurons appear to be relatively unstable circadian oscillators *in vivo*, exhibiting a stochastic pattern of rhythmicity in calcium dynamics over time. However, correlational network analysis reveals a strong, stable circadian pattern of coherence among AVP neuron pairs at the population level, peaking during the circadian daytime. This study is the first to undertake cell-type-specific *in vivo* recordings of individual SCN neurons, demonstrating that AVP neuronal rhythms emerge largely at the network level.

## Methods

### Animals

For *in vivo* calcium imaging experiments, 10-week-old heterozygous male AVP-IRES2-Cre-D (Jackson Laboratory 023530)[44] were used. Cre-mediated recombination in these mice specifically targets SCN AVP+ neurons in several studies to date [45, 46]. Control experiments were conducted using male Ai6 mice which express robust ZsGreen1 fluorescence following Cre-mediated recombination (Jackson Labs 007906) crossed with AVP-IRES2-CRE-D knock-in mice (Jackson Labs 023530). For *in vivo* optogenetically targeted single-unit recording, Mice were bred by crossing AVP-Cre with Ai32 (RCL-ChR2(H134R)/EYFP,Stock # 024109) mice. *In vivo* electrophysiological recordings were performed on adult male mice (3-4 months) housed in 12L:12D light-dark cycle from birth. Food and water were available *ad libitum*. All procedures described were approved by the IACUC committee at Morehouse School of Medicine.

### In vivo Miniaturized Fluorescent Calcium Imaging

Chronic lens implantation surgeries were done as previously described[47]. Briefly, using a stereotaxic apparatus, AVP-cre mice were injected (ML: +0.750 mm, AP: −0.200 mm, DV: −5.8 mm) with an adeno-associated viral vector conveying the genetically encoded calcium sensor jGCaMP7s (#104491, Addgene) into the SCN and a gradient index lens 0.50 mm in diameter (1050-004611 Inscopix) was implanted above the SCN (ML: +0.750 mm, AP: −0.200 mm, DV: −5.7 mm; **Figure 1A**). Minimitter G2 Emitters were implanted at the same time into the peritoneal cavity. Post-hoc histology verified lens placement near/above the SCN and away from other AVP-expressing cell populations in hypothalamic paraventricular nucleus, or supraoptic nucleus. This placement was targeted since GRIN lens focal distance is typically ~200 μm below the bottom surface. Fluorescence in ventral hypothalamus, but outside the SCN borders appeared non-cellular in nature, while cell bodies are clearly observable within the SCN. Six weeks after surgery, mice were assessed for evidence of discrete, dynamic calcium signals and a baseplate was attached over the gradient index lens to establish a fixed working distance with the microscope. Baseplated mice (n=4 mice) were acclimated to the isolated recording chamber and the presence of the microscope for 2 days before recordings were initiated. The recording schedule consisted of 5-minute recordings where mice were imaged at 6.67 Hz, every three hours for 48 consecutive hours for a total of 16 timepoints under both constant darkness and 12:12LD. An example field-of-view and recording traces (5 min) from 2 different ROIs and a background region containing no visible cell are shown in **Figure 1B**. The two 48-hour recordings did not result in photobleaching or any other run-down of GCaMP in the neurons. Specificity of GCaMP infection was verified by colocalization of GCaMP and AVP via IHC (**Figure 1C**). Colocalization was difficult to detect, as expected due to the reduced expression of the AVP peptide in AVP-cre mice[48]. GCaMP7s never colocalized with neurons expressing VIP, supporting AVP-expressing neuronal specificity. Locomotor activity (LMA) during a fixed, then a shifted LD cycle was measured in one mouse after imaging to verify that implanted mice had unimpaired circadian rhythms, including proper entrainment to phase shifts (**Figure 1D**). Coincident recording of LMA was also used to verify rhythmicity during Ca^2+^ recordings (**Figure S1**). Proper functionality of the calcium sensor was established by comparing dynamic GCaMP7s recordings with GFP expressing AVP neurons used as a control (**Figure 1E**). The GFP signal never exhibiting dynamics like those observed using the calcium indicator. **Figures 1F-G** show 3 more example traces to highlight the features of the traces used for the analysis described in more detail below.

**Figure 1:**
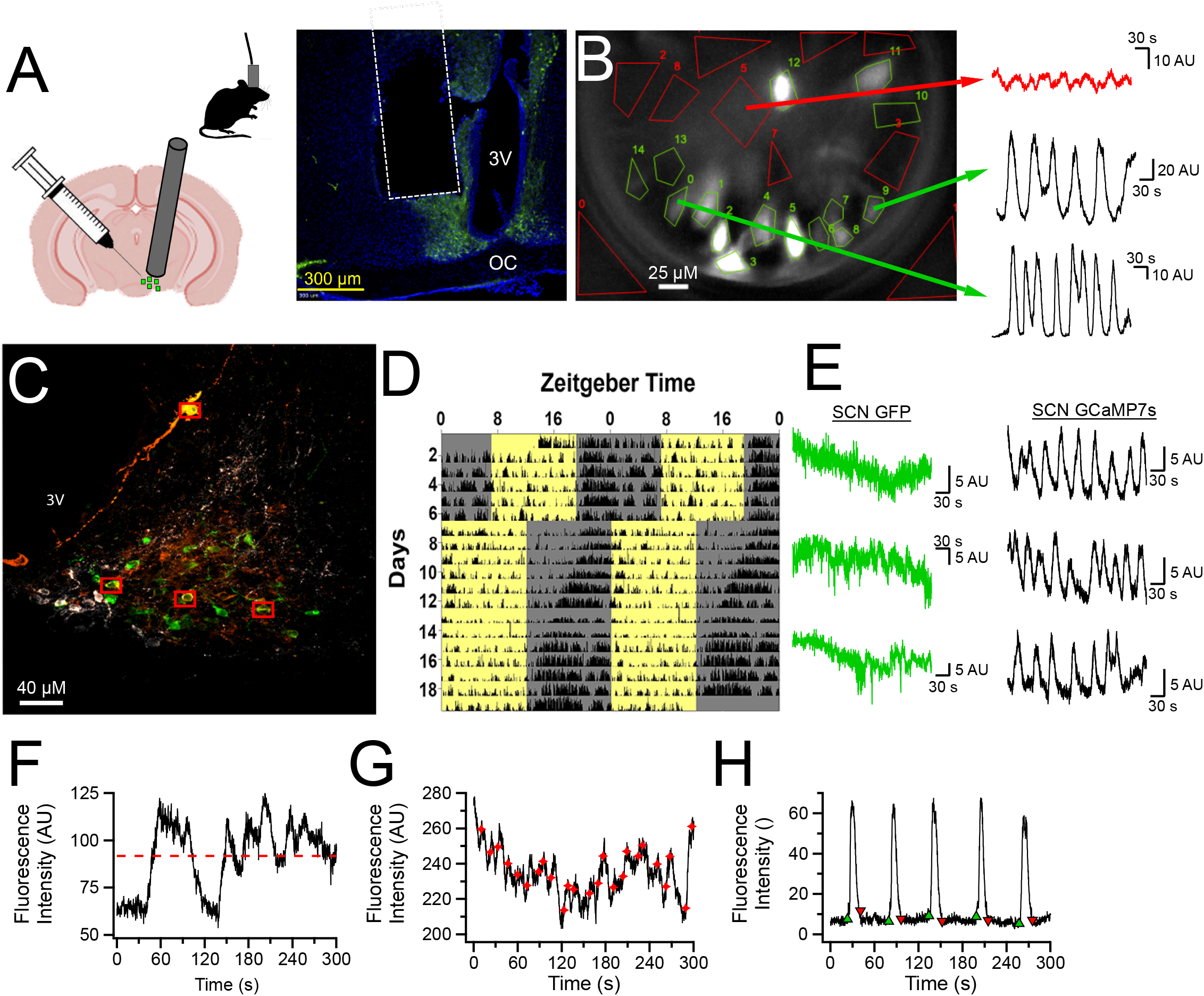
*In Vivo* Characterization of AVP Neurons in the SCN Network. **A**: Left: Illustrations describing AAV infection of AVP neurons, GRIN implantation above the SCN, and tethered recordings using microendoscope. Right: example of histological validation of lens placement. Typical focal distance below the lens is ~200um. **B**: Representative still image demonstrating how AVP calcium dynamics are observed *in vivo*. Green polygons are drawn around AVP neuronal regions of interest (ROIs) and the red polygons are non-AVP neuronal spaces used for background subtraction. Sample traces on the right illustrate examples of 5-minute recordings obtained from AVP neurons in the same field of view. **C**: Image of the SCN taken with a confocal microscope (63x lens) showing neuronal colocalization (yellow) of GCaMP7s (green) and AVP antibody (orange) as well as the exclusion of GCaMP7s from VIP expressing neurons (white). Red boxes indicate neurons with clear colocalization. **D**: Representative actogram demonstrating that mice with a GRIN lens implanted above the SCN appear to have normal daily circadian behavior which shifts normally in response to a 6-hour phase advance. **E**: Representative calcium traces recorded from the SCN of AVPcre x GFP mice and AVPcre mice infected with GCaMP7s demonstrating that while GFP does not exhibit acute calcium dynamics, intracellular calcium dynamics are detected by GCaMP7s. **F-H**: Representative 5-minute calcium traces demonstrating how mean fluorescence intensity (red dotted line, **F**), acute events (red stars, **G**), and calcium waves (onsets green; offsets red, **H**) were quantified for the purposes of determination of single-cell rhythmicity in Fig. 2.

**Figure 2:**
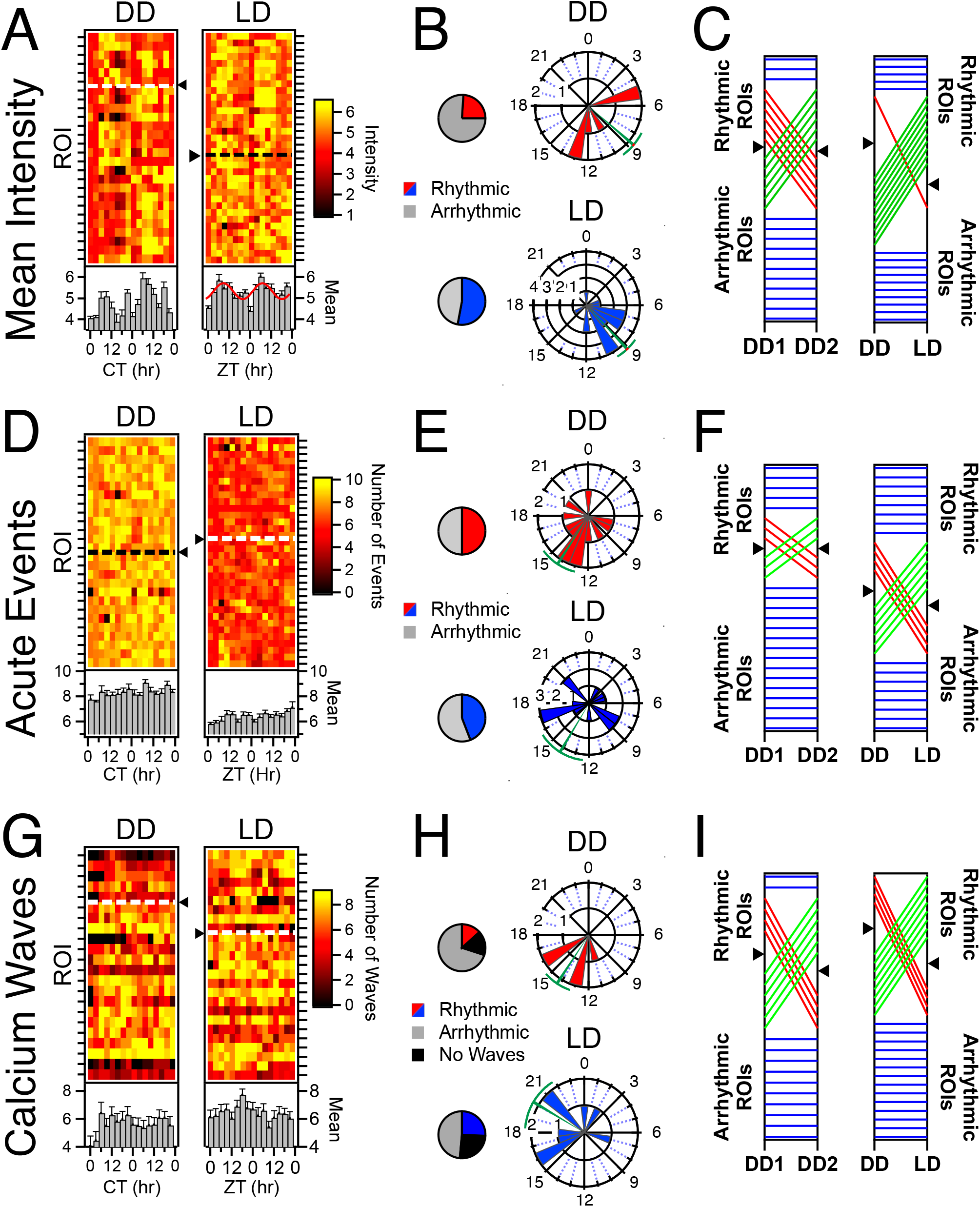
Circadian Rhythmicity in individual AVP Neurons is stochastic. **A**: Heatmaps illustrating mean fluorescence intensity across the 5-minute recording (see Fig. 1F) of individual AVP neurons/ROIs (rows) by recording timepoint (columns) in both DD and LD. Rows are normalized to their means. ROIs which exhibited significant circadian rhythms according to cosinor (p<0.05) for mean intensity appear above the dashed line and triangle. ROIs that were not rhythmic are plotted below the dashed line. Plotted beneath the heatmap is the population average and standard errors for each timepoint. When the population measure is rhythmic (p<0.05), the cosine fit is superimposed on top of the bar graph. **B**: Polar plots illustrating the phase distribution of ROIs exhibiting significant circadian rhythms in mean fluorescence in both DD and LD. The direction of the bar represents hour in CT for DD or ZT for LD. The length of the bar represents number of ROIs. Circular mean and SEM are represented in green. Pie charts indicate the proportion that exhibited a statistically significant rhythm across the 48 hours of DD and LD. **C**: Phenotype tracking plots indicating the stability/change of rhythmic state of individual AVP neuron ROIs between the 2 recording days (DD day 1 and DD day 2) under constant darkness (left) and between the 48h recordings in each lighting condition (right). Blue lines indicate stable rhythmic or arrhythmic ROIs, while red lines indicate a loss of rhythmicity between conditions, and green lines a gain of rhythmicity. Black triangles on the axes indicate the division between rhythmic (above) and arrhythmic (below) ROIs. **D-F**: Rhythmicity measures for unsupervised acute events as quantified as described in Fig. 1G. Same conventions as A-C, above, except that raw event number is plotted in the heatmaps rather than normalized counts. **E-I**: Rhythmicity measures for calcium waves, as quantified as described in Fig. 1H. Same conventions as A-C, above.

### In Vivo Optogenetically-targeted Single-unit Activity Recording (SUA^AVP^)

#### Optrode Implantation

Optrode Implantation was done as previously described[49]. Briefly, a stereotaxic apparatus (KOPF, model 1900) was used for implanting custom 12-channel optrodes above the SCN (AP: +0.38 mm, ML: +0.15 mm, DV: −5.5 mm). Optrodes were drivable via manual screw-drive, and once implanted were slowly advanced into the SCN over days/weeks (see below). Optrodes were constructed by combining a microelectrode bundle (insulated nichrome wire, 30μm diameter, Stablohm 675, California Fine Wire, USA) with a custom-made optical fiber (200 μm diameter, Doric Lenses). The entire site was then sealed and secured with dental acrylic (**Figure 3A**).

**Figure 3:**
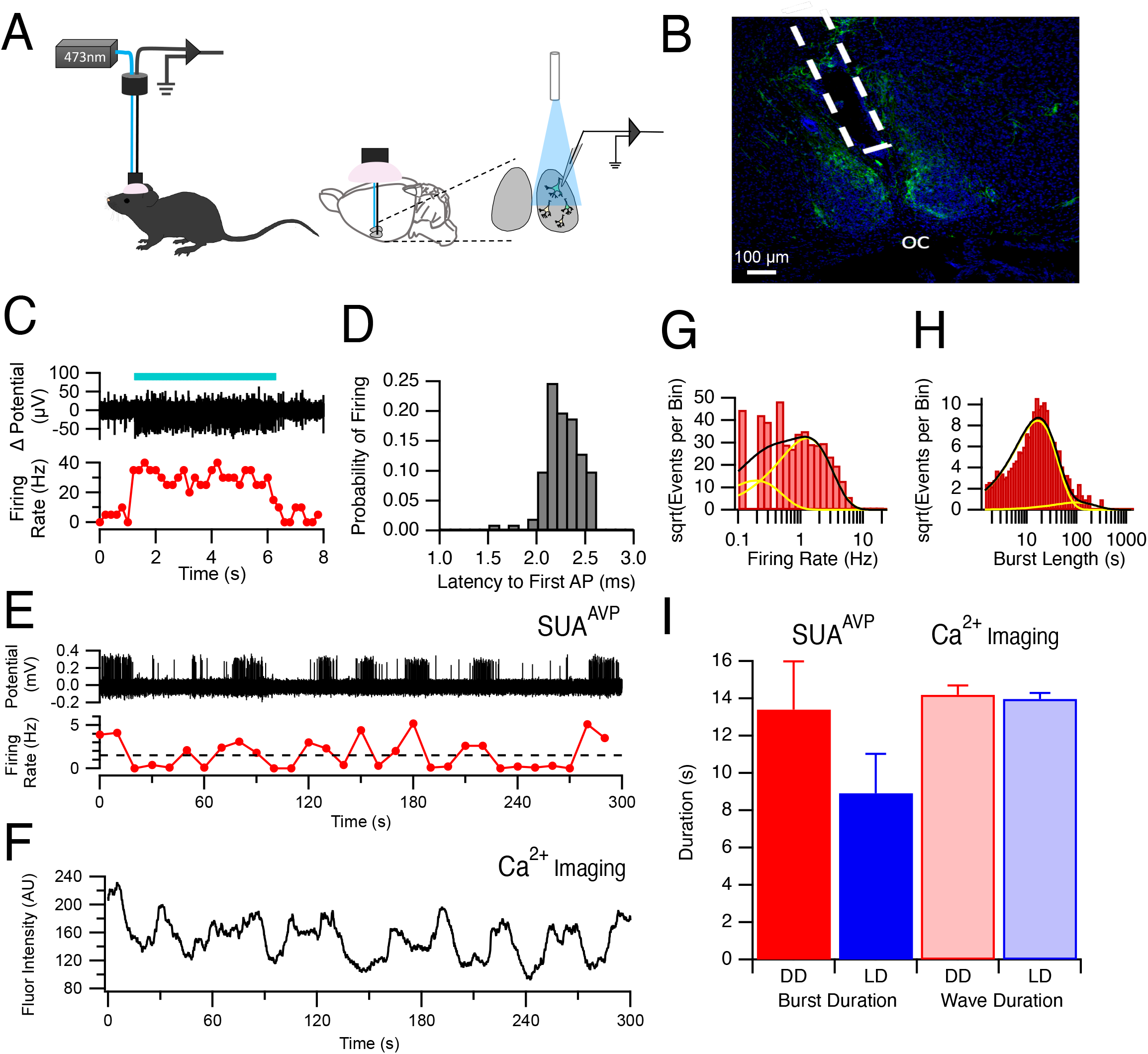
Electrical Characterization of Calcium Waves using Optogenetically-Tagged Single Unit Activity Recording *In Vivo*. **A**: Schematic illustrating how laser stimulation is used to identify and record from AVP neurons in freely behaving mice. **B**: Photomicrograph of representative histological section illustrating the location of the fiber-optrode above the SCN. Blue=DAPI; Green=EYFP expression tag for channelrhodopsin. OC=optic chiasm. **C**: Blue light stimulation (blue bar) yielded an increase in firing rate in AVP+ neurons containing channelrhodopsin. **D**: Histogram showing short (~2.25ms) latency between blue light stimulation and action potential firing in AVP neurons containing channelrhodopsin. **E**: Action potential firing measured from the optrode of a representative AVP neuron (top) with the smoothed firing rate plotted beneath. **F**: Representative 5-minute calcium trace of GCaMP mediated fluorescence from an AVP neuron. Note the similarity between the fluorescence calcium waves to the smoothed firing rate dynamics measured by SUA^AVP^ plotted in E. **G**: Example histogram of firing rate for an AVP neuron. The black line indicates the fit to the log transform of the exponential fit according to equation 3 with the yellow lines being the individual exponential components from the fit. **H**: Example histogram from the same neuron showing the log transform of burst length. **I**: Summary of average burst duration recorded with SUA^AVP^ (left) compared with the duration of calcium waves measured via miniaturized fluorescent microscopy (right). There was no significant difference in duration between recording modalities and lighting conditions (ANOVA p>0.05).

#### Identification of ChR2-Expressing Neurons and Data Acquisition

After recovery from electrode implantation surgery, the mice were handled to adapt to extracellular recording procedures (30 min) and the optrode is gradually lowered through the brain toward the SCN (40μm, once a day) until light stimulation through the optrode yielded single unit responses. For optogenetic identification of ChR2-expressing (AVP+) neurons, pulses of blue light (473 nm, 2-10 ms duration, 20 Hz, 10-15 mW) were delivered to the SCN through the optical fiber. A single unit was identified as a ChR2-expressing neuron when action potential spikes were repeatedly and reliably evoked by the blue light pulses (>50% occurrence) with a short first-spike latency (<7 ms), low jitter (<3 ms) and with waveforms consistent with prior-recorded spontaneous spikes (correlation coefficient between waveforms of spontaneous and optically evoked spikes >97%). The latency to action potential firing after light stimulation was 2.26 ms ±
0.02 ms (n = 100 randomly selected APs; **Figure 3D**). Thirteen AVP neurons were recorded from 5 mice in total. However, for only 3 of them (AVP neuron #3, 4, 13), recorded from 2 mice, were we able to obtain complete recordings of a successive 96 hours (48h under LD and 48h under DD). One neuron (AVP neuron #9) from one mouse was lost during recording under DD. So 4 neurons in LD and 3 neurons in DD were used for further data analysis. Electrophysiological data were recorded with a sampling rate of 20 KHz by a 16-channel head-stage and controller (RHD2000, Intan Technologies). Recorded raw voltage data was high-pass filtered (>250 Hz) and spikes were identified by applying thresholding. Single units were identified through offline sorting and then analyzed using NeuroExplorer and MATLAB software. All electrode locations were verified histologically (**Figure 3B**). Due to the relative scarcity of the neurons recorded, longitudinal (i.e. circadian) analysis was not performed for SUA^AVP^ data and instead the data were only used to analyze bursting behavior.

### Image Analysis and Statistics

A complete description of our data analysis approach is provided in **Supplemental Methods**. A brief description follows.

#### Image Correction, ROI Definition and Production of Intensity Traces

Raw image stacks acquired at 6.67 Hz were motion-corrected using Inscopix Data Processing software and five-minute recordings of images were exported as TIFF stacks. The TIFF stacks were imported into Igor Pro 8 (Wavemetrics Inc, Lake Oswego, Oregon) for further analysis. Regions of interest (ROIs) were identified from maximum projection images. Using the background subtracted image stack, the average intensity for each ROI was determined for each image in the stack. This produced a line trace of varying intensities with time over the five-minute period for each ROI (e.g., **Figure 1F-H**).

#### Mean Intensity, Intensity Correlation, and Events Analysis

To determine whether individual ROIs varied with circadian rhythmicity, we first characterized each ROI for its average intensity within the five-minute trace at each 3-hour time point (**Figure 1F, Figure 2A-C**). Fluorescence intensity traces for each ROI were also cross-correlated with each other yielding a Pearson coefficient for each ROI-ROI interaction at each circadian time point (**Figure 4A**). In addition, we performed event analysis on the calcium traces. This was accomplished in two different ways. An unbiased analysis of acute events was conducted in which the onset of an event was found if the amplitude increased by more than two standard deviations above the intensity of the previous 20 points (3 seconds) (**Figure 1G, Figure 2 D-F**). A second method was used to determine event parameters of slow calcium waves (**Figure 1H, Figure 2 G-I**). In a first pass, automatic analysis was done in which the raw fluorescence line trace for each ROI was smoothed with a 50-point box window. Then, the first derivative was calculated from the smoothed raw data trace. The first derivative trace was also smoothed using a 100-point box window followed by recording the time when a threshold level on the first derivative was crossed in an increasing manner when searching from the start of the first derivative trace to the end for either positive or negative slopes. This approximated the start of slow calcium fluorescence wave rise and the end of the calcium wave. The value of the threshold of the first derivative was determined heuristically (usually 0.05). These time points could then be manually corrected for missing and/or spurious events. Wave durations, inter-event intervals and wave number could then be calculated from the start and end times of each event in each fluorescent intensity line trace.

**Figure 4:**
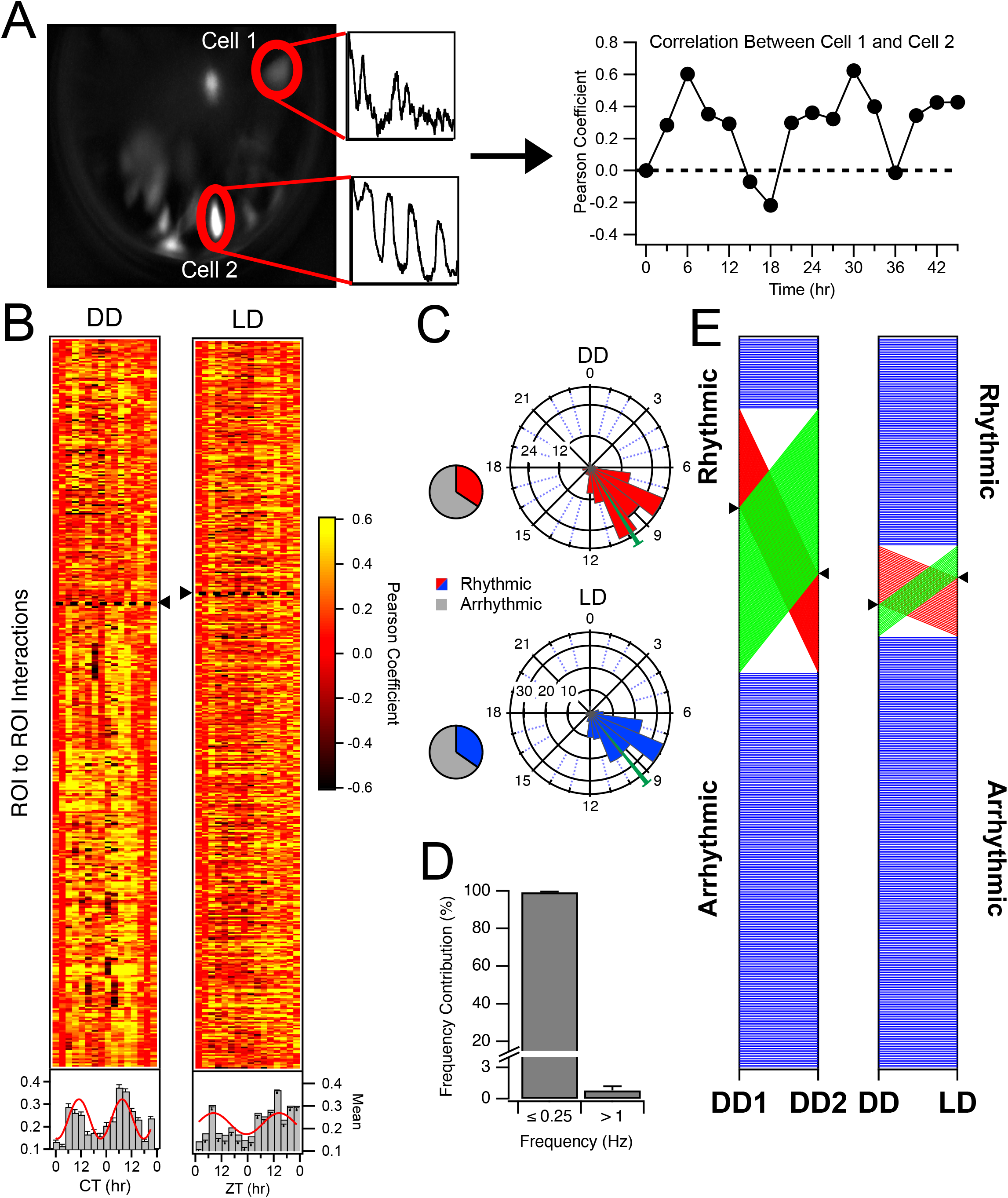
Circadian Rhythmicity in Correlation Among AVP Neuron Pairs. **A**: Schematic illustrating correlational (a.k.a. coherence) analysis among AVP neuron pairs. Calcium traces of individual SCN neurons across a field-of-view within a 5-minute recording were cross-correlated, then the Pearson’s coefficient for cell pairs was plotted for each time point over 48 hours and subjected to rhythmicity analysis. High correlation is interpreted as active interaction or coordination among cell pairs and referred to as *coherence*. Coherence among some pairs was strongly rhythmic. **B**: Heat maps illustrating Pearson coefficients of single AVP neuronal pairs (rows) by timepoint (columns) in both DD and LD. Neuronal pairs which exhibited significant circadian rhythms according to cosinor (p<0.05) in correlated activity appear above the dashed line and triangle. Plotted beneath the heatmap is the population average and standard errors for each timepoint. Where the population measure is rhythmic (or trending in that direction), the cosine fit is superimposed on top of the bar graph (DD cosinor p=0.004; LD cosinor p=0.057). **C**: Polar plots illustrating the phase distribution of ROI pairs exhibiting significant circadian rhythms in coherence in both DD and LD. The direction of the bar represents hour in CT for DD or ZT for LD. The length of the bar represents number of ROI pairs. Pie charts indicate the proportion that exhibited a statistically significant rhythm across the 48 hours of DD and LD. **D**: Histogram illustrating the frequency contribution to neuronal pair correlation. A low-pass filter (<0.25Hz) applied to raw data yielded correlations with Pearson Coefficients approximating the unfiltered data. On average, over 90% of the power was carried by these low frequencies in comparison to when a high-pass filter (>1 Hz) was applied to the raw data. **E**: Phenotype tracking plots indicating the stability/instability of rhythmic state of AVP neuron ROI pair coherence between the 2 recording days (DD day 1 and DD day 2) under constant darkness (left) and between the 48h recordings in each lighting condition (right). Blue lines indicate stable rhythmic or arrhythmic pairs, while red lines indicate a loss of rhythmicity between conditions, and green lines a gain of rhythmicity. Black triangles on the axes indicate the division between rhythmic (above) and arrhythmic (below) ROI pairs.

#### Circadian Rhythm Analysis

Circadian Rhythmicity was tested by fitting data (mean intensity, acute events, calcium wave variables including number, duration and inter-event interval, and Pearson correlations among ROI pairs) collected at three-hour intervals over 24 or 48 hours with a cosine function. Fit optimization was based on Levenberg-Marquardt least-squares method constraining the results for the period between 21 and 28 hours[50, 51]. The *p*-values from these fits were determined using a non-parametric Mann-Kendall tau test. A particular ROI was deemed rhythmic if the *p*-value was below 0.05. The 24- or 48-hour data was then ranked by *p*-value and displayed as heatmaps. Data which could not be fit with Equation 2 was given a *p*-value of 1 in the resulting heatmaps and is not ranked (**Figures 2A, 2D, 2G, 4B**). Those non-rhythmic ROIs are plotted below the dashed line in each heatmap.

To determine if AVP neurons displayed any rhythmicity as a population, the columns of the heatmaps were averaged, and these values were subjected to the circadian fit. For population averages, fitting was also weighted by the reciprocal of the standard deviations determined for each column of the heatmap. When the population measure was rhythmic, the cosine fit is displayed on the mean bar graph below each heat map (**Figures 2A, 2D, 2G, 4B**).

Phases resulting from the fits to Equation 2 were recorded for each ROI or ROI-ROI interaction with significant rhythmicity. Histograms of the phases were then calculated utilizing 3-hour bins and then plotted on polar coordinate graphs (**Figures 2B, 2E, 2H and 4C**) with associated circular means and SEM (in green).

#### Calculation of Firing Rate and Duration of Bursting for Single Unit Analysis

After spike sorting, traces dedicated to a particular single unit were analyzed for their firing rate (**Figure 3E**). Time stamps of action potential firing were subjected to histogram analysis with a bin size of 3 s over the whole recording and then divided by the bin size to get the firing rate per 3 s time point. A histogram of this firing rate time course was then produced, indicating the prevalence of firing rates. Burst lengths were determined by rebinning the time stamp data per 10 s bins and determining the duration of the burst by analyzing the points when the firing rate had risen above and then returned below 1.5 Hz (**Figure 3G**). Because firing rates and burst lengths could have a range of several decades, these histograms were converted to a log transform where the abscissa was the log of the firing rate and the ordinate was the square root of the number of events per bin (e.g. **Figures 3G, 3H**)[52]. This data was then fit with a log transformation of a sum of exponential components [53] to determine the firing rate or the burst length for **Figures 3G** and **3H**, respectively.

Raw data from this study can be found at https://doi.org/10.5061/dryad.2ngf1vhpz.

## Results

### Circadian Rhythmicity in Mean Fluorescence of AVP Neurons in the SCN

Circadian analysis of mean fluorescence intensities could indicate if basal levels of calcium, a surrogate for overall levels of neuronal activity, would change over the course of the day. Average fluorescence intensity was determined from each 5-minute recording at each three-hour time point for each ROI (e.g. **Figure 1F**). The mean fluorescence per three-hour time point was fit with equation 2 for each ROI to determine if the ROI exhibited a circadian rhythm. Heatmaps of these rhythms in mean intensity are shown in **Figure 2A**. In constant darkness, 24% of AVP neurons exhibited significant circadian rhythmicity (cosinor, p<0.05) with mean peak phase at CT 8.69 ± 0.92 hours; n=6 ROIs (**Figure 2A, left, above the dotted line; Figure 2B upper)**; whereas in 12:12 LD, 53% of the AVP neurons demonstrated significant circadian rhythmicity with a mean peak phase at ZT 9.11 ± 0.80 hours; n=18 ROIs (**Figure 2A, right, above the dotted line; Figure 2B lower**).

For each individual 24-hour day, approximately 35-50% of recorded cells were rhythmic, but the set of cells that made up that rhythmic subpopulation was dynamic, with neurons gaining or losing rhythmicity in the two 24-hour periods (**Figure 2C left; Figure S4A)**. Also, there was no discernable pattern between lighting conditions except for an overall increase in the likelihood of rhythmicity in LD compared with DD (**Figure 2C right**). When mean fluorescence values for all ROIs were averaged for each 3-hour time point, the population did not display circadian rhythmicity in constant darkness (**Figure 2A; cosinor p = 0.546; n=26 ROIs**). In contrast, under a 12:12 LD cycle the population average resulted in a significant population rhythm in mean fluorescence (**Figure 2B; cosinor p = 0.005; n=34 ROIs)**, peaking during late day.

### Rhythmicity of Acute Calcium Events in AVP Neurons

Although slower variants of GCaMP including the GCaMP7s used in this study are not capable of resolving distinct rapid events such as action potentials, significant changes in fluorescence within a recording are indicative of neuronal dynamics in these AVP neurons. Within the 5-minute recordings, events were detected automatically by finding fluorescence that crossed a threshold that was two standard deviations above a sliding 3-second window (e.g. red diamonds, **Figure 1G**). In DD, 50% of AVP neurons exhibited a circadian rhythm (cosinor p<0.05) in the number of dynamic calcium events per 5-minute recording session compared with 44.1% of neurons in LD (**Figure 2D)**. The sub-population average phase for these dynamic events for these rhythmic cells in DD was at CT 14.21 ± 1.30 hours (n=13 ROIs) with a similar population average phase for LD at ZT 14.10 ± 1.68 hours (n=15 ROIs) (**Figure 2E**).However, when considering all characterized AVP neurons (both rhythmic and non-rhythmic), the population average exhibited no significant rhythm in DD (**Figure 2D, left below the heatmap;** cosinor p = 0.322; n=26 ROIs) or LD (cosinor p = 0.652; n=34 ROIs). This was largely due to the wide phase distribution apparent in the polar plots for the rhythmic ROIs (**Figure 2E**). As was observed for mean fluorescence (**Figure 2C**), there was a fairly consistent ~30% of neurons that were rhythmic for any given 24-hour period, but the members of that rhythmic subset, and sometimes the phase of those rhythms (**Figure S4H**), were highly dynamic across the 2 recording days of DD (**Figure 2F, left**) and LD (**Figure S4B**), and between 48-hour series from each lighting conditions (**Figure 2F, right**).

### AVP Neurons in the SCN Exhibit Distinctive Calcium Waves in vivo

Slow, high amplitude calcium waves appear to be a characteristic feature of most AVP neurons in the SCN. Nearly all ROIs in three of the four mice recorded for this study exhibited unambiguous waves distinct from the other dynamics apparent in the signal (See **Table S1**). For the 4^th^ mouse (AVP63), the waves were less distinct from the faster dynamics. Such waves were never observed in control experiments in which AVP neurons expressing GFP under the CAG promoter were recorded using the same methods, in either DD or LD (**Figure 1E**). The waves occurring within each 5-minute recording were counted (**Figure 2H**) across the circadian day to determine if the frequency of these events is rhythmic. To be conservative, wave counting and analysis for rhythmicity was not performed for AVP63 (see **Table S1**). In DD, calcium waves were observed in most AVP neurons during at least 1 (usually most) time point(s), though a significant circadian rhythm in their counts was observed in only 16.7% of AVP neurons with a peak phase at CT 14.24 ± 0.96 hours; n=5 ROIs (**Figure 2G, Figure 2H**). Under 12:12 LD, calcium waves were also observed in most analyzed neurons, and significant circadian rhythms were detectable in 26.5% of the cells with a peak phase at ZT 20.06 ± 1.79 hours; n=9 (**Figure 2G, Figure 2H**). Averaging the number of waves by timepoint for all AVP neurons did not reveal a significant circadian population rhythm in DD or in LD (**Figure 2G below the heatmaps;** p = 1 (n=26 ROIs); p = 0.133 (n=34 ROIs) respectively). As reported above for both mean fluorescence and acute calcium events, very little stability in the subpopulation of rhythmic ROIs was observed for the number of calcium waves over 2 days in DD (**Figure 2I, left)** or LD (**Figure S4C**), with a variable ~15-30% of neurons rhythmic for 24-hour periods. And again, for calcium waves as reported above for other measures, largely different populations of cells were rhythmic in DD or LD (**Figure 2I, right**). In addition to the analysis of wave number here, wave duration (**Figure S2**) and inter-event interval between these calcium waves (**Figure S3**) were also calculated. Neither additional measure provided evidence of stable circadian rhythmicity or common phasing, and both exhibited similar state-switching (**Figures S2D, S3D, S4, S5**) as reported above.

### Calcium Waves Likely Correspond to Bursting Activity in AVP Neurons

Optogenetically-targeted Single Unit Activity Recording (SUA^AVP^) was employed to determine if the timing of the slow wave phenomenon observed in calcium imaging (e.g. **Figure 1H**) corresponded to increased action potential firing activity in individual AVP neurons within the intact SCN (**Figures 3A, 3B**). Mice expressing channelrhodopsin in AVP neurons were implanted with optrodes and neurons were verified as AVP positive if an increase in firing rate was elicited by blue light stimulation through the optrode (**Figure 3C**). The latency to action potential firing after light stimulation was 2.26 ms ± 0.02 ms (n = 100 randomly selected Aps; **Figure 3D**). When the firing rate was considered for 5-minute epochs, a distinct pattern of bursting was observed (**Figure 3E**) which corresponded closely with the calcium imaging data observed from SCN AVP neurons in other animals (**Figure 3F**). Both firing rate (**Figure 3G**) and burst duration (**Figure 3H**) were calculated using the unbiased method described above. Average burst length was 13.41 ± 2.6 seconds and 8.92 ± 2.11 seconds in DD (n=3 cells) and LD (n=4 cells) respectively compared with the calcium wave duration of 14.20 ± 0.50 seconds and 13.97 ± 0.32 seconds in DD and LD respectively (**Figure 3I**). Two-way ANOVA indicated that there was no main effect of method of observation (DF = 1, F_1,677_ = 1.27, p = 0.26; n =7) or lighting condition (DF = 1, F_1,677_ = 0.827, p = 0.37, n=7) as well as no interaction (DF = 1, F_1,677_ = 0.67, p = 04.1, n=7).

### Rhythmicity of Correlated Activity between AVP Neurons

Raw fluorescence across all cell/ROI pairs within a recording (**Figure 4A**) was subjected to correlational analysis for each time-point to quantify the degree of coherence,or coordinated activity among AVP neuron pairs. We sought to determine if the degree of coherence varies over circadian and diurnal time, which would indicate changes in the state of network signaling or output beyond those appreciated by single-neuron analysis, or by measurement of bulk fluorescence. We observed that many ROI pairs exhibited strong positive correlations between their calcium dynamics during the middle/late day/subjective day and either no correlation or were inversely correlated during the subjective night (**Figure 4A, Right**). In DD, 34.33% of AVP neuron pairs exhibited a circadian rhythm in correlated activity (**Figure 4B left; ROI pairs above the dotted line)** with an average peak correlation for rhythmic pairs at a phase of CT 9.83 ± 0.25 hours; n=149 ROI pairs (**Figure 4C, upper**). In LD, 34.70% of ROI pairs demonstrated significant rhythmicity of coherence with a mean peak phase at ZT 9.37 ± 0..24 hours; n=161 ROI pairs (**Figure 4B right; Figure 4C lower**). In both cases, the phase distributions were tight compared with measures of single cell rhythms in Figure 2. The population average of Pearson coefficients by timepoint exhibited a significant rhythm at the population level (all cells/ROIs) in DD and in LD (**Figure 4B, below the heatmap;** DD: p=0.004; n=434 ROI pairs; LD: p = 0.024; n=451 ROI pairs). Most ROIs in the study had at least 1 rhythmic correlation with another ROI, with the modal # of rhythmic connections being from 2-5 (**Figure S6C**).

We wanted to know how much of the power in the correlations resulted from low frequency fluorescence changes (< 0.25 Hz) or if higher frequency calcium dynamics were mostly responsible for the coherence between AVP neuron ROIs. Frequency contribution analysis on a subset of interactions indicated that wave components slower than 0.25 Hz accounted for 99.2 ± 0.40% of the correlation between AVP neurons (n=10 ROI pairs; **Figure 4D)**, indicating that an increased synchronous activity was arising from coincidence in calcium waves or other slow dynamics in AVP neuronal pairs rather than due to shared high-frequency dynamics.

Rhythmicity in correlated activity consistently occurred in ~25-35% of AVP neuron pairs across all four of the 24-hour periods for which we recorded (**Figure 4E, S4**), but the membership of that rhythmic subset of cell pairs was highly variable. Cell pairs either gained, lost, or maintained their rhythmic relationship across the 48-hour recordings (**Figure 4E left, Figure S4**) and between DD and LD (**Figure 4E right**) with no predictability. The strength of correlation as a function of distance between neurons was also measured. In DD, there was a weak but significant relationship between correlated activity and the distance between AVP neurons, such that the further one neuron was from another for both positively (R^2^ = 0.007, p<0.001; n=160 ROI pairs) and negatively correlated relationships (R^2^ = 0.01, p<0.004;n=160 ROI pairs), Pearson coefficients decreased. Correlated activity also weakly decreased with distance between neurons for 12:12LD, for both positively correlated (R^2^ = 0.02, p<0.001;n=188 ROI pairs) and negatively correlated (R^2^ = 0.04, p<0.001;n=188 ROI pairs).

### Multimodal analysis of rhythmicity

**Table S1** provides a summary of rhythmic characteristics across all of our primary measures for each ROI. The final two columns indicate how many rhythmic coherence relationships each ROI has within that mouse. Likelihood of rhythmicity across variables appears to be stochastic, such that no one parameter predicts the value of another, and no single-cell variable predicts the number of rhythmic relationships. Results from each mouse in the study share these general characteristics. **Figure S6** summarizes the multimodal analyses with histograms that indicate shared and unique rhythmic features within ROIs. About 60% of cells exhibited at least one rhythmic single-cell parameter in both DD and LD (Figure S6A and S6B), and nearly all cells had at least one rhythmic relationship, with the modal # of rhythmic relationships of 2-4.

## Discussion

### Individual neuron heterogeneity of function in the context of an intact circuit

The master circadian pacemaker in mammals in the suprachiasmatic nucleus (SCN) is comprised of groups of neurons which express specific neuropeptides and these cells may exert distinct influences over the regulation of circadian rhythms. It is still poorly understood how this network generates a unified output in which different neurons within the network interact to regulate circadian rhythmicity. Imaging of brain slices, using bioluminescent and fluorescent gene-reporter mice, demonstrates the existence of robust network organization that is predictably and reversibly altered by changing light conditions[11–13, 54]. However, this approach is limited by the need to dissect the SCN network for observation, severing all inputs, outputs, and most intra-network connections. Multi-unit recording offers a window into the electrical activity of the intact SCN network *in vivo*, and such studies have been fundamental in developing our understanding of the SCN as a master pacemaker for circadian rhythmicity [28–31, 55], but are unable to provide additional information regarding cell-type-specific population or individual neuronal contribution to the regulation of circadian rhythmicity within the SCN.

Intersectional genetics and novel recording techniques have dramatically improved our ability to conduct population level, cell-type specific studies within the intact circadian network. For example, recent studies *in vivo* demonstrate the necessity and sufficiency of VIP-expressing neurons in synchronizing the SCN to light-induced phase resetting[23, 32, 33]. *In vivo* fiber photometry is a powerful technique, but one that can be enhanced with the ability to observe individual neurons and the interactions between these neurons within the fully intact SCN network. Thus, here we present work involving two novel techniques: optogenetically-targeted single unit activity recording (SUA^AVP^) and *in vivo* calcium imaging of AVP-expressing neurons in the intact SCN of awake, freely moving mice under different lighting schedules using miniaturized fluorescence microscopy. Targeted SUA has been used for cell-type specific electrophysiological recording in the medial prefrontal cortex[56], and the striatum[57], but applied here for the first time in the SCN. *In vivo* calcium imaging has been used to measure calcium dynamics in many brain areas including the frontal cortex[58], hippocampus[59], and cerebellum[60, 61]. This technique permits the characterization of individual cells within the SCN and how they contribute to communal network behavior, and how these contributions are modulated over circadian and diurnal time.

Our data indicate that most AVP neurons in the SCN exhibit slow calcium waves, approximately 20% of which show a modulatory rhythm of their frequency over a 24-hour period (**Figure 2G-I**). While increases in intracellular calcium can be indicative of neuronal firing [62], because of the slow decay time of the variant of GCaMP7 we employed, temporal discrimination of individual action potentials within bursting events was not possible. Thus, we employed SUA^AVP^ to better characterize the firing behavior of AVP neurons within the SCN *in vivo*. The burst duration resulting from firing rate of AVP neurons by SUA^AVP^ is similar to the duration of calcium waves measured by fluorescence (**Figure 3I**). This suggests that the waves we observe in our calcium imaging data are likely the result of bursting activity exhibited by AVP neurons. Previous studies observed similar bursting activity in AVP neurons of the supraoptic nucleus[63] and paraventricular nucleus[64]. Although the specific cell-types were not identified, such bursting was also reported within the SCN[65]. The present results are the first to bridge these findings and confirm that AVP neurons within the SCN exhibit phasic firing, or bursting activity, *in vivo*. Interestingly, although bursting activity is present in nearly all SCN AVP neurons recorded, a minority of neurons exhibit circadian rhythmicity in bursting. This suggests that while bursting is an identifying characteristic of these neurons, it may not play a significant role in the regulation of circadian rhythms.

Although circadian or diurnal regulation of bursting is uncommon in SCN AVP+ neurons (**Figures 2G-I, S2, S3**), the well-known role of these cells as regulators of the circadian clock suggests that some aspect of their activity should exhibit reliable rhythmicity. AVP-specific *Per2∷luc* rhythms in SCN slices illustrate the presence of functional molecular clocks[26]. We have observed similar results using a D site-binding protein (DBP) reporter in AVP-expressing SCN cells [42], however our selection of cell-based regions-of-interest used rhythmicity as a criterion and thus excluded non-rhythmic cells which may have been present. Imaging from SCN slices from AVPEluc/+ mice demonstrates the rhythmic transcription of *avp* in the SCN[66], and there is a circadian rhythm in the secretion of vasopressin[67, 68] and GABA[34] from AVP neurons in the SCN.

Using longitudinal calcium imaging, we find that stable circadian rhythmicity is not observed in all AVP neurons. Only a minority of these neurons exhibited rhythmicity for each of the 3 main parameters characterized (mean fluorescent intensity, calcium events, or calcium waves). While individual neurons sometimes exhibit circadian rhythmicity in various measures (sometimes several measures; **Table S1** and **Figure S6**), these rhythms are not stable from day to day nor is there a reliable trend in rhythmicity between lighting conditions (**Figures 2, S2, S3, S4**). Furthermore, the phases of the calcium rhythms that we did observe vary among cells as well as across time and lighting condition such that rhythms do not emerge at the population level in some of these measures. This was not equally true among the 3 measures, as mean intensity was rhythmic in LD at the population level, and was trending that way in DD. But overall, while AVP neurons exhibit strong rhythms in gene transcription with coherent phasing in slice recordings, such rhythmicity may not accurately reflect the temporal, and electrical activity of these neurons *in vivo*. Instead, we suggest that stable rhythmic activity of individual neurons is not needed. Instead, perhaps a generous (and ever-changing) subset (i.e., ~30% of cells perhaps) need only be rhythmic, electrically, in order to drive rhythmic outputs. These data may suggest a novel view of network rhythmicity for AVP neurons, where all cells contribute to the generation of a population rhythm, but not all the time. A very similar phenomenon has been reported for hippocampus, where the dynamic sharing of the responsibility for encoding place memory among neurons was described [69].

At first blush, this view of SCN rhythmic organization may seem at-odds with what’s been reported prior, that a large majority of SCN neurons exhibit rhythms in gene expression and bulk Ca^2+^ which reflect underlying molecular rhythms in these cells [7, 8, 11, 40]. We argue that our findings reflect properties only appreciable through longitudinal *in vivo* recordings with cellular resolution. It is important to distinguish how these cells might behave in the presence of dynamic input from outside of the SCN from how they behave in a slice. It is also important to distinguish how a cell behaves over time, versus how the population does. Previous studies utilizing diverse techniques have had either <u>single-cell resolution</u> in a reduced prep, OR <u>*in vivo*</u> bulk recordings, OR *in vivo* single cell recordings without cellular specificity. No studies have had all of these advantages combined, and thus could not have observed the stochasticity and dynamically shared function reported here. The closest comparison is *in vivo* MUA/SUA [70, 71], which (until now) lacks cell specificity, but nonetheless has revealed non-rhythmic cells, cells with diverse phases, and acute cellular behavior which reflects dynamic behavioral input, which are observations one would predict given our current results. Thus, the suggestion of diversity among neurons and stochastic behavior of rhythms of individual neurons in the SCN circuit *in vivo* is not surprising. Such diversity of rhythmicity [43] and phase [40] both among cells and over time within cells has been reported for SCN. Phasic diversity among cells *in vivo* may also play an important role in plasticity and robustness of the circuit [41]. Recordings of other brain circuits using the same technique employed in our study suggests that heterogeneity of function, even among cells that share many other features is the norm, not the exception[35–38], and is likely a fundamental property of neural circuits.

### Correlated activity among cells during only the daytime is an emergent network property of AVP neurons

Recent work suggests that emergent properties of the SCN network play an important role in the generation of robust and stable rhythmicity. Emergent properties are observations that cannot be appreciated by studying single cells alone, and that may have functional significance for how the network operates. The wave observed in bioluminescent slice recordings reveal a spatiotemporal organization within the SCN [11, 14, 15] which is predictably and reversibly altered by changes in environmental lighting[16, 17]. Synchronization between cells may be dependent on this organization[72, 73] driven by region-specific circadian programming within the SCN network[54, 74, 75]. However, to date it is unknown if such an organization observed *ex vivo* extends to an intact SCN network *in vivo*. Previous *in vivo* studies were not capable of single-cell discrimination and/or have not been able to observe multiple cells simultaneously, and therefore have not addressed the issue of network organization. Here, while single-cell rhythms appear unstable, we observed an underlying circadian structure to coordinated activity among neuron pairs. Calcium dynamics in many AVP cell pairs are more highly correlated (i.e. *coherent*) during the day than during the night, indicating either coordinated function intrinsic to those cells, or a shared acute response to a common input of either intrinsic or extrinsic origin (**Figure 4**). This coherence in activity is dependent on distance such that the further two neurons are from each other in the same field of view, the less correlated they are likely to be. Generally, the relationship is absent during the night such that the cells behaved entirely independently, or sometimes are inversely correlated. About 33% of all AVP neuron pair relationships have significant circadian rhythms, and nearly every cell we recorded has at least one such relationship (most have more) (**Figure S6C**). The average of these correlations across all cell pairs, a surrogate population measure, in both DD and LD exhibit a significant circadian rhythm with a (subjective) daytime peak between CT/ZT 6-9. This *in vivo* observation of rhythms in coherence may reflect time-of-day dependent signaling dynamics in the SCN previously reported. GABA is known to switch between being inhibitory and excitatory in a time-of-day dependent manner [76–79], regulated via neuronal intracellular chloride concentration and expression of the Na^+^-K^+^-2Cl^−^ cotransporter (NKCC1). The significance of this is demonstrated by time-dependent blockade of the NKCC1 transport, which results in diminished phase-resetting to light exposure during the early subjective night, but not during the late subjective night or during the light phase[80]. Changing network dynamics may also be driven thru non-synaptic paracrine signaling (e.g. by AVP itself). The relevance of this time-of-day dependent rhythm in neuronal coherence for circadian function needs further study.

### Study Limitations

There are technical limitations to this body of work that place some constraints upon the conclusions^.^ Due to the difficulty in cell-type specific targeting of a relatively small neuronal population (~2,000 neurons) within a very small, deep brain region, the number of AVP neurons which were observed in a longitudinal fashion was somewhat limited. Now that feasibility of this technique has been established by us and others [81], other cell specific populations can be examined. The AVP-IRES2-Cre mouse used in this study has been reported to express reduced levels of AVP protein and mRNA, with no change in circadian behavior or AVP cell number [48]. This fact limited our ability to demonstrate robust colocalization of AVP and gcAMP7 expression using immunofluorescence. However the model is well-documented to selectively target AVP neurons in the SCN and elsewhere [42, 44–46]. Additionally, there is the possibility that a minority of AVP-expressing neurons dorsal to, but not part of, the SCN that are marked by the cre-line may have been recorded [82]. It was not possible to distinguish between these and AVP-expressing neurons within the SCN in the present study. Since the vast majority of AVP-expressing neurons in the ventromedial hypothalamus are within the SCN [19], it is almost certain that most cells recorded are within its borders. No lens was found to be dorsal enough to have recorded from the hypothalamic paraventricular nucleus, and lateral placements that captured supraoptic neurons were unambiguous and omitted from this analysis. In **Figure 1A**, the fluorescence that is present dorsal to the SCN is largely non-cellular in appearance. And since the heterogeneity we report is longitudinal across time in addition to across cells, ectopic AVP-neurons present in our recordings would not account for such diversity. Future studies will take advantage of novel cell position identification techniques such as light-guided sectioning [83] to identify precisely where, and what, recorded neurons are *post-hoc*, allowing us to determine whether they exhibit different behavior inside versus outside the SCN.

Another potential limitation is that damage caused by implantation of the GRIN lens could have altered the function of the SCN. Tissue damage is also a limitation for other techniques such as slice recording (neo-natal or adult), *in vivo* MUA and fiber photometry approaches. The question being addressed with SUA^AVP^ in the present study did not require recording from many neurons, resulting in a relatively low number of observations. This precluded us from doing an accurate longitudinal analysis. Future studies will include full 48-hour observations of a larger number of cells in both in DD and LD using this parallel technique. Finally, since the changing behavioral state of animals may modify cellular function acutely, behavioral and sleep measures will be added to future cellular imaging experiments to assess this possibility and to better understand the functions of these neurons within the context of behavior.

The population measure of rhythmic coherence among AVP cell pairs highlights the importance of emergent characteristics in the SCN neural circuit as well as the potential limitations of studying population level dynamics which ignore individual neuronal activity. While the use of cell-type specific AVP recording in this study helped to ensure we were observing network activity mostly within the SCN, the relative sparsity of this neuronal subtype made observing a large number of neurons prohibitive. Furthermore, conducting such analysis on only a single cell-type within a heterogenous network is inherently limiting. Therefore, future studies will focus on investigating the role of network organization in a more diverse, SCN-specific cell population. Given what we have now learned about AVP cell behavior *in vivo*, recording a broader cohort of cell types inclusive of, but extending beyond just AVP neurons, will reveal whether our findings are reflective of behavior unique to AVP cells, or reflect general properties of SCN neuronal function.

## Acknowledgements

This work was supported by National Institute of Health grants R21NS108197 and R35GM136661 to AJD, SC1AG046907 to JCE, NHLBI-funded predoctoral fellowship to DMB (T32HL007901) and an equipment grant to MSM by the W.M. Keck Foundation. We also thank Ivory Ellis and Inscopix, Inc., for technical assistance with the project.

The authors declare no conflicts of interest.

---

**Supplemental Table S1. Summary of rhythmic and non-rhythmic parameters by ROI and mouse.** All ROIs analyzed for this study across 4 mice are arrayed in rows. Checkmarks indicate the detection of statistically significant circadian rhythmicity (Cosinor fit p<0.05) for each of our major measures, for 48h DD and LD datasets. The final 2 columns list the # of rhythmic coherence relationships each ROI had with others from the same mouse. Note that for AVP10, a DD recording was incomplete and not included in the dataset. Discrete, unambiguous Ca^2+^ waves were less common in AVP63, so waves were not counted for rhythmicity for this mouse.

**Supplemental Figure S1. Behavioral rhythmicity. All 4 mice recorded from for this study had Minimitter G2 emitters implanted.** Robust rhythmicity of locomotor behavior was verified for each mouse prior to recordings. Shown here is an example of locomotor activity (Mouse AVP40) collected contemporaneously with the Ca2+ recordings used for analysis. Raw behavior actograms are shown for 48h DD, 48h LD. Below are the associated periodograms for each 48h recording.

**Supplemental Figure S2. Circadian Rhythmicity in Duration of Dynamic Calcium Events of Individual AVP Neurons. A**: Representative 5-minute calcium trace demonstrating analysis of duration of calcium waves (time between the green (onset) and red (offset) triangles). **B**: Heat maps illustrating the duration of calcium waves of individual AVP neurons (rows) by timepoint (columns) in both DD and LD. Neurons which exhibited significant circadian rhythms in calcium wave duration appear above the dashed line. Plotted beneath are the population averages and standard errors for each timepoint. For DD the population was arrhythmic according to cosinor, while in DD a rhythm was detected (p=0.013; cosine superimposed on histogram). **C**: Polar plots illustrating the phase distribution, and pie charts showing the proportion of AVP neurons/ROIs exhibiting significant circadian rhythms in calcium waves duration in both DD and LD. The direction of the bar represents hour in CT for DD or ZT for LD. The length of the bar represents number of ROIs. Mean phase ± SEM is indicated in green. **D**: Phenotype tracking plots indicating the stability/change in rhythm state of individual AVP ROIs between recording days (DD Day1-DD Day2) under constant darkness (left) and between lighting conditions (right). Red lines indicate a loss of rhythmicity between conditions, green lines a gain of rhythmicity, and blue lines indicate no change in rhythm state between conditions. Black triangles on the axes indicate the division between rhythmic (above) and arrhythmic (below) ROIs.

**Supplemental Figure S3. Circadian Rhythmicity in Inter-event Interval of Dynamic Calcium Events of Individual AVP Neurons. A**: Representative 5-minute calcium trace demonstrating analysis of calcium wave inter-event interval (time between the red (offset) triangle of one wave and the green (onset) triangles of the next wave). **B**: Heat maps illustrating the inter-event interval of calcium waves of individual AVP neurons (rows) by timepoint (columns) in both DD and LD. Neurons which exhibited significant circadian rhythms in inter-event interval of calcium waves appear above the dashed line and triangle. Plotted beneath are the population averages and standard errors for each timepoint. Population-level circadian rhythmicity was not observed for DD or LD (cosinor p> 0.05). **C**: Polar plots illustrating the phase distributions, and pie charts the proportion of AVP neurons/ROIs exhibiting significant circadian rhythms in inter-event interval of calcium waves in both DD and LD. The direction of the bar represents hour in CT for DD or ZT for LD. The length of the bar represents number of ROIs. Mean phase ± SEM is indicated in green. **D**: Phenotype tracking plots indicating the stability/change in rhythm state of individual AVP ROIs between recording days (DD Day 1-DD Day 2) under constant darkness (left) and between lighting conditions (right). Red lines indicate a loss of rhythmicity between conditions, green lines a gain of rhythmicity, and blue lines indicate no change in rhythm state between conditions. Black triangles on the axes indicate the division between rhythmic (above) and arrhythmic (below) ROIs.

**Supplemental Figure S4. Additional single-day measures of rhythmicity. A-F**: Phenotype tracking plots indicating the stability/change in rhythmic state of individual AVP ROIs (A-E), or pairwise correlated relationships (F) between recording days in a LD cycle (LD Day 1-LD Day 2). Red lines indicate a loss of rhythmicity between conditions, green lines a gain of rhythmicity, and blue lines indicate no change in rhythm state between conditions. Black triangles on the axes indicate the division between rhythmic (above) and arrhythmic (below) ROIs. **G-I**: Polar plots showing phase of rhythms in ROIs for individual days in the study across the 3 primary measures of single-cell rhythms.

**Supplemental Figure S5. Additional single-day measures of rhythmicity. A-C**: Single-day polar plots for correlations (A), wave duration (B) and wave inter-event interval (C). Same conventions as previous figures.

**Supplemental Figure S6. Multimodal summary of rhythmicity across measures. A-B**: Histograms showing how rhythmicity in various single-cell measures overlaps within ROIs in 48h constant darkness (A) and in a 48h LD cycle (B). Furthest to the right, the blue histograms indicate the fraction of total ROIs that were rhythmic in *at least* 1, 2 or all 3 major parameters. **C**: Histogram reporting the number of pairwise rhythmic correlations per ROI for 48hDD (C) and 48hLD(D) datasets. Note that some mouse recordings had more ROIs than others (See Table S1), making a higher number of rhythmic correlations possible.

## Cited References

1. Ralph, M.R., et al., Transplanted suprachiasmatic nucleus determines circadian period. Science, 1990. 247(4945): p. 975–8.

2. Weaver, D.R., The suprachiasmatic nucleus: a 25-year retrospective. J Biol Rhythms, 1998. 13(2): p. 100–12.

3. Takahashi, J.S., Transcriptional architecture of the mammalian circadian clock. Nat Rev Genet, 2017. 18(3): p. 164–179.

4. Welsh, D.K., et al., Individual neurons dissociated from rat suprachiasmatic nucleus express independently phased circadian firing rhythms. Neuron, 1995. 14(4): p. 697–706.

5. Su Terman, J., C.E. Rem′, and M. Terman, Rod outer segment disk shedding in rats with lesions of the suprachiasmatic nucleus. Brain Research, 1993. 605(2): p. 256–264.

6. Koronowski, K.B., et al., Defining the Independence of the Liver Circadian Clock. Cell, 2019. 177(6): p. 1448–1462.e14.

7. Yoo, S.-H., et al., PERIOD2∷LUCIFERASE real-time reporting of circadian dynamics reveals persistent circadian oscillations in mouse peripheral tissues. Proceedings of the National Academy of Sciences of the United States of America, 2004. 101(15): p. 5339–5346.

8. Herzog, E.D., et al., Circadian rhythms in mouse suprachiasmatic nucleus explants on multimicroelectrode plates. Brain Research, 1997. 757(2): p. 285–290.

9. Honma, S., et al., Circadian periods of single suprachiasmatic neurons in rats. Neuroscience Letters, 1998. 250(3): p. 157–160.

10. Herzog, E.D., J.S. Takahashi, and G.D. Block, Clock controls circadian period in isolated suprachiasmatic nucleus neurons. Nature Neuroscience, 1998. 1(8): p. 708–713.

11. Noguchi, T., et al., Calcium Circadian Rhythmicity in the Suprachiasmatic Nucleus: Cell Autonomy and Network Modulation. eNeuro, 2017. 4(4).

12. Schmal, C., E.D. Herzog, and H. Herzel, Measuring Relative Coupling Strength in Circadian Systems. Journal of Biological Rhythms, 2018. 33(1): p. 84–98.

13. Yamaguchi, S., et al., Synchronization of Cellular Clocks in the Suprachiasmatic Nucleus. Science, 2003. 302(5649): p. 1408.

14. Hastings, M.H., et al., Differential regulation of mPER1 and mTIM proteins in the mouse suprachiasmatic nuclei: new insights into a core clock mechanism. J Neurosci, 1999. 19(12): p. Rc11.

15. Enoki, R., et al., Topological specificity and hierarchical network of the circadian calcium rhythm in the suprachiasmatic nucleus. Proceedings of the National Academy of Sciences, 2012. 109(52): p. 21498.

16. Evans, J.A., et al., Dynamic interactions mediated by nonredundant signaling mechanisms couple circadian clock neurons. Neuron, 2013. 80(4): p. 973–83.

17. Rohr, K.E., et al., Seasonal plasticity in GABA(A) signaling is necessary for restoring phase synchrony in the master circadian clock network. Elife, 2019. 8.

18. Moore, R.Y. and J.C. Speh, GABA is the principal neurotransmitter of the circadian system. Neurosci Lett, 1993. 150(1): p. 112–6.

19. Abrahamson, E.E. and R.Y. Moore, Suprachiasmatic nucleus in the mouse: retinal innervation, intrinsic organization and efferent projections. Brain Res, 2001. 916(1–2): p. 172–91.

20. Meng, Q.J., et al., Setting clock speed in mammals: the CK1 epsilon tau mutation in mice accelerates circadian pacemakers by selectively destabilizing PERIOD proteins. Neuron, 2008. 58(1): p. 78–88.

21. Lee, I.T., et al., Neuromedin s-producing neurons act as essential pacemakers in the suprachiasmatic nucleus to couple clock neurons and dictate circadian rhythms. Neuron, 2015. 85(5): p. 1086–102.

22. Park, J., et al., Single-Cell Transcriptional Analysis Reveals Novel Neuronal Phenotypes and Interaction Networks Involved in the Central Circadian Clock. Front Neurosci, 2016. 10: p. 481.

23. Jones, J.R., et al., SCN VIP Neurons Are Essential for Normal Light-Mediated Resetting of the Circadian System. The Journal of Neuroscience, 2018. 38(37): p. 7986.

24. Mieda, M., et al., Cellular Clocks in AVP Neurons of the SCN Are Critical for Interneuronal Coupling Regulating Circadian Behavior Rhythm. Neuron, 2015. 85(5): p. 1103–1116.

25. Mieda, M., H. Okamoto, and T. Sakurai, Manipulating the Cellular Circadian Period of Arginine Vasopressin Neurons Alters the Behavioral Circadian Period. Current Biology, 2016. 26(18): p. 2535–2542.

26. Shan, Y., et al., Dual-Color Single-Cell Imaging of the Suprachiasmatic Nucleus Reveals a Circadian Role in Network Synchrony. Neuron, 2020. 108(1): p. 164–179.e7.

27. Todd, W.D., et al., Suprachiasmatic VIP neurons are required for normal circadian rhythmicity and comprised of molecularly distinct subpopulations. Nature Communications, 2020. 11(1): p. 4410.

28. Inouye, S.T. and H. Kawamura, Persistence of circadian rhythmicity in a mammalian hypothalamic “island” containing the suprachiasmatic nucleus. Proc Natl Acad Sci U S A, 1979. 76(11): p. 5962–6.

29. Yamazaki, S., et al., Rhythmic Properties of the Hamster Suprachiasmatic Nucleus*In Vivo*</em&gt. The Journal of Neuroscience, 1998. 18(24): p. 10709.

30. Nakamura, W., et al., In vivo monitoring of circadian timing in freely moving mice. Curr Biol, 2008. 18(5): p. 381–5.

31. Takasu, N.N., et al., In vivo monitoring of multi-unit neural activity in the suprachiasmatic nucleus reveals robust circadian rhythms in Period1^−^/^−^ mice. PLoS One, 2013. 8(5): p. e64333.

32. Ono, D., K. Honma, and S. Honma, Circadian and ultradian rhythms of clock gene expression in the suprachiasmatic nucleus of freely moving mice. Sci Rep, 2015. 5: p. 12310.

33. Mazuski, C., et al., Entrainment of Circadian Rhythms Depends on Firing Rates and Neuropeptide Release of VIP SCN Neurons. Neuron, 2018. 99(3): p. 555–563.e5.

34. Maejima, T., et al., GABA from vasopressin neurons regulates the time at which suprachiasmatic nucleus molecular clocks enable circadian behavior. Proceedings of the National Academy of Sciences, 2021. 118(6): p. e2010168118.

35. Jarvie, B.C., et al., Satb2 neurons in the parabrachial nucleus mediate taste perception. Nat Commun, 2021. 12(1): p. 224.

36. Jennings, J.H., et al., Visualizing hypothalamic network dynamics for appetitive and consummatory behaviors. Cell, 2015. 160(3): p. 516–27.

37. Viskaitis, P., et al., Modulation of SF1 Neuron Activity Coordinately Regulates Both Feeding Behavior and Associated Emotional States. Cell Rep, 2017. 21(12): p. 3559–3572.

38. Zimmerman, C.A., et al., A gut-to-brain signal of fluid osmolarity controls thirst satiation. Nature, 2019. 568(7750): p. 98–102.

39. Bellesi, M., et al., Enhancement of sleep slow waves: underlying mechanisms and practical consequences. Front Syst Neurosci, 2014. 8: p. 208.

40. Evans, J.A., et al., Intrinsic regulation of spatiotemporal organization within the suprachiasmatic nucleus. PLoS One, 2011. 6(1): p. e15869.

41. Evans, J.A., et al., Neural correlates of individual differences in circadian behaviour. Proc Biol Sci, 2015. 282(1810).

42. Smith, C.B., et al., Cell-Type-Specific Circadian Bioluminescence Rhythms in Dbp Reporter Mice. J Biol Rhythms, 2022. 37(1): p. 53–77.

43. Webb, A.B., et al., Intrinsic, nondeterministic circadian rhythm generation in identified mammalian neurons. Proc Natl Acad Sci U S A, 2009. 106(38): p. 16493–8.

44. Pei, H., et al., AVP neurons in the paraventricular nucleus of the hypothalamus regulate feeding. Mol Metab, 2014. 3(2): p. 209–15.

45. Shan, Y., et al., Dual-Color Single-Cell Imaging of the Suprachiasmatic Nucleus Reveals a Circadian Role in Network Synchrony. Neuron, 2020. 108(1): p. 164–179 e7.

46. Whylings, J., et al., Reduction in vasopressin cells in the suprachiasmatic nucleus in mice increases anxiety and alters fluid intake. Horm Behav, 2021. 133: p. 104997.

47. Resendez, S.L., et al., Visualization of cortical, subcortical and deep brain neural circuit dynamics during naturalistic mammalian behavior with head-mounted microscopes and chronically implanted lenses. Nat Protoc, 2016. 11(3): p. 566–97.

48. Cheng, A.H., S.W. Fung, and H.M. Cheng, Limitations of the Avp-IRES2-Cre (JAX #023530) and Vip-IRES-Cre (JAX #010908) Models for Chronobiological Investigations. J Biol Rhythms, 2019. 34(6): p. 634–644.

49. Cardin, J.A., et al., Targeted optogenetic stimulation and recording of neurons in vivo using cell-type-specific expression of Channelrhodopsin-2. Nature Protocols, 2010. 5(2): p. 247–254.

50. PES volume 2 issue 1 Cover and Back matter. Probability in the Engineering and Informational Sciences, 1988. 2(1): p. b1–b3.

51. Shrager, R.I., Quadratic programming for nonlinear regression. Commun. ACM, 1972. 15(1): p. 41–45.

52. Sigworth, F. and S. Sine, Data transformations for improved display and fitting of single-channel dwell time histograms. Biophysical journal, 1987. 52(6): p. 1047–1054.

53. Colquhoun, D. and F. Sigworth, Fitting and statistical analysis of single-channel records, in Single-channel recording. 1995, Springer. p. 483–587.

54. Enoki, R., et al., Dual origins of the intracellular circadian calcium rhythm in the suprachiasmatic nucleus. Scientific Reports, 2017. 7(1): p. 41733.

55. Michel, S., et al., Electrophysiological Approaches to Studying the Suprachiasmatic Nucleus. Methods Mol Biol, 2021. 2130: p. 303–324.

56. Anikeeva, P., et al., Optetrode: a multichannel readout for optogenetic control in freely moving mice. Nature Neuroscience, 2012. 15(1): p. 163–170.

57. Kravitz, A.V., S.F. Owen, and A.C. Kreitzer, Optogenetic identification of striatal projection neuron subtypes during in vivo recordings. Brain Research, 2013. 1511: p. 21–32.

58. Prevedel, R., et al., Fast volumetric calcium imaging across multiple cortical layers using sculpted light. Nature Methods, 2016. 13(12): p. 1021–1028.

59. Indersmitten, T., et al., Utilizing Miniature Fluorescence Microscopy to Image Hippocampal Place Cell Ensemble Function in Thy1.GCaMP6f Transgenic Mice. Current Protocols in Pharmacology, 2018. 82(1): p. e42.

60. Sullivan, M.R., et al., In Vivo Calcium Imaging of Circuit Activity in Cerebellar Cortex. Journal of Neurophysiology, 2005. 94(2): p. 1636–1644.

61. Díez-García, J., W. Akemann, and T. Knöpfel, In vivo calcium imaging from genetically specified target cells in mouse cerebellum. NeuroImage, 2007. 34(3): p. 859–869.

62. Fleming, W., et al., Inferring spikes from calcium imaging in dopamine neurons. PLOS ONE, 2021. 16(6): p. e0252345.

63. Gouzènes, L., et al., Vasopressin Regularizes the Phasic Firing Pattern of Rat Hypothalamic Magnocellular Vasopressin Neurons. The Journal of Neuroscience, 1998. 18(5): p. 1879.

64. Dewald, M., et al., Phasic bursting activity of paraventricular neurons is modulated by temperature and angiotensin II. Journal of Thermal Biology, 1999. 24(5): p. 339–345.

65. Miller, J.D. and C.A. Fuller, Isoperiodic neuronal activity in suprachiasmatic nucleus of the rat. American Journal of Physiology-Regulatory, Integrative and Comparative Physiology, 1992. 263(1): p. R51–R58.

66. Yoshikawa, T., et al., Spatiotemporal profiles of arginine vasopressin transcription in cultured suprachiasmatic nucleus. European Journal of Neuroscience, 2015. 42(9): p. 2678–2689.

67. Södersten, P., et al., A daily rhythm in behavioral vasopressin sensitivity and brain vasopressin concentrations. Neuroscience Letters, 1985. 58(1): p. 37–41.

68. Kalsbeek, A., et al., In vivo measurement of a diurnal variation in vasopressin release in the rat suprachiasmatic nucleus. Brain Research, 1995. 682(1): p. 75–82.

69. Ziv, Y., et al., Long-term dynamics of CA1 hippocampal place codes. Nat Neurosci, 2013. 16(3): p. 264–6.

70. Meijer, J.H., et al., Multiunit activity recordings in the suprachiasmatic nuclei: in vivo versus in vitro models. Brain Res, 1997. 753(2): p. 322–7.

71. Schaap, J., et al., Heterogeneity of rhythmic suprachiasmatic nucleus neurons: Implications for circadian waveform and photoperiodic encoding. Proc Natl Acad Sci U S A, 2003. 100(26): p. 15994–9.

72. Paszek, P., et al., Population robustness arising from cellular heterogeneity. Proceedings of the National Academy of Sciences, 2010. 107(25): p. 11644.

73. Gu, C. and H. Yang, Differences in intrinsic amplitudes of neuronal oscillators improve synchronization in the suprachiasmatic nucleus. Chaos: An Interdisciplinary Journal of Nonlinear Science, 2017. 27(9): p. 093108.

74. Enoki, R., et al., Synchronous circadian voltage rhythms with asynchronous calcium rhythms in the suprachiasmatic nucleus. Proceedings of the National Academy of Sciences, 2017. 114(12): p. E2476.

75. Pauls, S., et al., Differential contributions of intra-cellular and inter-cellular mechanisms to the spatial and temporal architecture of the suprachiasmatic nucleus circadian circuitry in wild-type, cryptochrome-null and vasoactive intestinal peptide receptor 2-null mutant mice. European Journal of Neuroscience, 2014. 40(3): p. 2528–2540.

76. Wagner, S., et al., GABA in the mammalian suprachiasmatic nucleus and its role in diurnal rhythmicity. Nature, 1997. 387(6633): p. 598–603.

77. Choi, H.J., et al., Excitatory Actions of GABA in the Suprachiasmatic Nucleus. The Journal of Neuroscience, 2008. 28(21): p. 5450–5459.

78. Jeu, M.D. and C. Pennartz, Circadian Modulation of GABA Function in the Rat Suprachiasmatic Nucleus: Excitatory Effects During the Night Phase. Journal of Neurophysiology, 2002. 87(2): p. 834–844.

79. Irwin, R.P. and C.N. Allen, GABAergic signaling induces divergent neuronal Ca2+ responses in the suprachiasmatic nucleus network. European Journal of Neuroscience, 2009. 30(8): p. 1462–1475.

80. McNeill, J.K., J.C. Walton, and H.E. Albers, Functional Significance of the Excitatory Effects of GABA in the Suprachiasmatic Nucleus. Journal of Biological Rhythms, 2018. 33(4): p. 376–387.

81. Cheikh Hussein, L.E., et al., Nested calcium dynamics support daily cell unity and diversity in the suprachiasmatic nuclei of free-behaving mice. bioRxiv, 2021: p. 2021.12.14.472553.

82. Rohr, K.E., et al., Vasopressin regulates daily rhythms and circadian clock circuits in a manner influenced by sex. Horm Behav, 2021. 127: p. 104888.

83. Kahan, A., et al., Light-guided sectioning for precise *in situ* localization and tissue interface analysis for brain-implanted optical fibers and GRIN lenses. Cell Reports, 2021. 36(13). https://doi.org/10.5061/dryad.2ngf1vhpz https://datadryad.org/stash/share/XZRVhWmI7Cd1X1mzvzTSoXAEys8IyfbUDg1ykL8jt9Q

