## Supplemental Methods for "Arginine-Vasopressin Expressing Neurons in the Murine Suprachiasmatic Nucleus Exhibit a Circadian Rhythm in Network Coherence *In Vivo*"

*Image Analysis and Statistics*

Image Correction, ROI Definition and Production of Intensity Traces

Raw image stacks acquired at 6.67 Hz were motion-corrected using Inscopix Data Processing software and five-minute recordings of images were exported as TIFF stacks. The TIFF stacks were imported into Igor Pro 8 (Wavemetrics Inc, Lake Oswego, Oregon) for further analysis. For each TIFF stack, a maximum projection image was then produced by finding the maximum intensity for each pixel during the 5-minute recording. This maximum projection image was used to identify regions of interest (ROIs) and regions of background. For a particular animal, ROI locations could vary slightly over the 24 – 48 hours of experimentation, and thus, ROI locations could be slightly corrected from 3-hour time point to time point. Background was subtracted from each image in the stack by doing a two-dimensional interpolation of background intensities based on the background regions outlined in the maximum projection image. Using the background subtracted image stack, the average intensity for each ROI was determined for each image in the stack. This produced a line trace of varying intensities with time over the five-minute period for each ROI (e.g., **Figure 1F-H**).

Mean Intensity, Intensity Correlation and Events Analysis

One focus of this study was to determine if activity of individual neurons varied with circadian rhythmicity. Thus, we characterized each ROI for its average intensity within the five-minute trace at each 3-hour time point (**Figure 1F, Figure 2A-C**). Fluorescence intensity traces for each ROI were also cross-correlated with each other yielding a Pearson coefficient for each ROI-ROI interaction at each circadian time point (**Figure 4A**). We wished to determine if the power in the cross-correlations mainly resulted from slow frequencies below 1 Hz or faster frequencies. To do this, we subjected raw data to high or low pass digital Finite Impulse Response filters before cross-correlation analysis. Parameters for the low pass filter were as follows- End Band Pass: 0.25 Hz; Start of Stop Band: 0.5 Hz; Number of Computed Terms: 73. Parameters of high pass filtering were as follows- End of First Band: 0.5 Hz; Start of Second Band: 1 Hz; Number of terms: 41. Edge effects were removed by deleting the first 17 and last 17 points in the filtered data trace. After cross-correlation analysis, the resulting Pearson Coefficients were analyzed by determining the weights that each of correlation of the filtered traces would contribute to the correlation coefficient that was not digitally filtered:

$P_{unfiltered}=w_{low pass}P_{low pass}+w_{high pass}P_{high pass}$	[1]

where *P* represents the Pearson Coefficient determined for unfiltered data or for the Pearson Coefficient after filtering the data with either high or low pass filters. *w* represents the contributing weights of each Pearson Coefficient to the Pearson Coefficient determined for the unfiltered data. An additional constraint was that:

*w_low pass_* + *w_high pass_* = 1

In addition, we performed event analysis on the calcium traces. This was accomplished in two different ways. An unbiased analysis of acute events was conducted in which the onset of an event was found if the amplitude increased by more than two standard deviations above the intensity of the previous 20 points (3 seconds) (**Figure 1G, Figure 2 D-F**). Data was smoothed prior to this analysis using a 30-point box window. A second method was used to determine event parameters of slow calcium waves (**Figure 1H, Figure 2 G-I**). In a first pass, automatic analysis was done in which the raw fluorescence line trace for each ROI was smoothed with a 50-point box window. Then, the first derivative was calculated from the smoothed raw data trace. The first derivative trace was also smoothed using a 100-point box window followed by recording the time when a threshold level on the first derivative was crossed in an increasing manner when searching from the start of the first derivative trace to the end for either positive or negative slopes. This approximated the start of slow calcium fluorescence wave rise and the end of the calcium wave. The value of the threshold of the first derivative was determined heuristically (usually 0.05). These time points could then be manually corrected for missing and/or spurious events. Wave durations, inter-event intervals and wave number could then be calculated from the start and end times of each event in each fluorescent intensity line trace.

Circadian Rhythm Analysis

Circadian Rhythmicity was tested by fitting data collected at three-hour intervals over 24 or 48 hours with a cosine function as follows:

$y=\frac{A}{2}\cos\left( \frac{2\pi\left( t-\psi\right)}{\theta} \right)+\frac{A}{2}+A_{o}$	[2]

where *A* represents the amplitude of the cosine signal, *A_o_* is the average offset of the fluorescence signal from zero, *t* is the time of each data point in hours, *ψ* and *θ* are the phase and period respectively in hours resulting from the fit. The data that were fit with Equation 2 to determine rhythmicity were the mean intensities, intensity correlations between ROIs, the wave number for dynamic calcium events as well as the number of calcium waves. Before fitting mean intensities, data was subjected to linear detrending to compensate for any consistent, progressive loss of fluorescence intensity over the period of the experiment. Fit optimization was performed using the internal Igor Pro algorithm based on Levenberg-Marquardt least-squares method constraining the results for the period between 20 and 32 hours. The *p*-values from these fits were determined using the Igor Pro implementation of the non-parametric Mann-Kendall tau test. A measure for a particular ROI was deemed rhythmic if the *p*-value was below 0.05. The 24- or 48-hour data was then ranked by *p*-value and displayed as heatmaps. Data which could not be fit with Equation 2 was given a *p*-value of 1 in the resulting heatmaps and is not ranked (**Figures 2A, 2D, 2G, 4B**). Those non-rhythmic ROIs are plotted below the dotted line in each heatmap.

To determine if AVP neurons displayed any rhythmicity as a population, the columns of the heatmaps were averaged, and these values subject to the circadian fit using Equation 2. For population averages, fitting was also weighted by the reciprocal of the standard deviations determined for each column of the heatmap. When the population measure was rhythmic, the cosine fit is displayed on the mean bar graph below each heat map (**Figures 2A, 2D, 2G, 4B**).

Phases (*ψ*) resulting from the fits to Equation 2 were saved for each ROI or ROI-ROI interaction with significant rhythmicity. Histograms of the phases were then calculated utilizing 3-hour bins and then plotted on polar coordinate graphs (**Figures 2B, 2E, 2H and 4C**).

Calculation of Firing Rate and Duration of Bursting for Single Unit Analysis

After spike sorting, traces dedicated to a particular single unit were analyzed for their firing rate (**Figure 3E**). Time stamps of action potential firing were subjected to histogram analysis with a bin size of 3 s for over the whole recording and then divided by the bin size to get the firing rate per 3 s time point. A histogram of this firing rate time course was then made, indicating the prevalence of firing rates. Burst lengths were determined by rebinning the time stamp data per 10 s bins and determining the duration of the burst by analyzing the points when the firing rate had risen above and then returned below 1.5 Hz (**Figure 3G**). 1.5 Hz was determined empirically, as it seemed to correlate with bursting behavior observed by eye. Because firing rates and burst lengths could have a range of several decades, these histograms were converted to a log transform where the abscissa was the log of the firing rate and the ordinate was the square root of the number of events per bin (e.g. **Figures 3G**, **3H**). This data was then fit with a log transformation of a sum of exponential components using the following equation[50]:

$y=A_{0}+\sum_{i}^{n} \left( \frac{A_{i}}{\iota_{i}}e^{\left( t-\frac{e^{t}}{\iota_{i}} \right)} \right)$	[3]

where *A*_i_ is the amplitude of the exponential of component *i*, *A*_0_ is the amplitude offset, *ι*_i_ is the exponential time constant of component *i* and n is the number of exponentials in the log transform. The variable *t* is the firing rate or the burst length for **Figures 3G** and **3H**, respectively.
