## Supplementary material for "Arginine-Vasopressin Expressing Neurons in the Murine Suprachiasmatic Nucleus Exhibit a Circadian Rhythm in Network Coherence *In Vivo*": Table S1

| **Summary of Circadian Rhythmicity** | | | | | | | | |
| --- | --- | --- | --- | --- | --- | --- | --- | --- |
| *Mouse ID - ROI #* | *Intensity* | | *Acute Events* | | *Ca Wave #* | | *# Rhythmic Correlations* | |
|  | **DD** | **LD** | **DD** | **LD** | **DD** | **LD** | **DD** | **LD** |
| **AVP10-0** | n/a |  | n/a |  | n/a |  | n/a | 3 |
| **AVP10-1** | n/a | **🗸** | n/a |  | n/a |  | n/a | 4 |
| **AVP10-2** | n/a | **🗸** | n/a | **🗸** | n/a |  | n/a | 5 |
| **AVP10-3** | n/a | **🗸** | n/a |  | n/a |  | n/a | 5 |
| **AVP10-4** | n/a |  | n/a |  | n/a |  | n/a | 5 |
| **AVP10-5** | n/a |  | n/a |  | n/a |  | n/a | 4 |
| **AVP10-6** | n/a |  | n/a |  | n/a | **🗸** | n/a | 6 |
| **AVP10-7** | n/a |  | n/a |  | n/a |  | n/a | 6 |
| **AVP24-0** |  | **🗸** |  | **🗸** |  |  | 5 | 5 |
| **AVP24-1** |  |  | **🗸** | **🗸** |  |  | 4 | 3 |
| **AVP24-2** |  |  |  |  |  |  | 6 | 2 |
| **AVP24-3** |  | **🗸** | **🗸** | **🗸** |  | **🗸** | 3 | 2 |
| **AVP24-4** |  |  | **🗸** |  |  | **🗸** | 3 | 2 |
| **AVP24-5** |  |  |  | **🗸** |  | **🗸** | 8 | 2 |
| **AVP24-6** | **🗸** |  | **🗸** | **🗸** |  |  | 3 | 6 |
| **AVP24-7** | **🗸** | **🗸** | **🗸** |  |  | **🗸** | 6 | 2 |
| **AVP24-8** |  | **🗸** |  | **🗸** |  |  | 4 | 7 |
| **AVP24-9** | **🗸** | **🗸** | **🗸** | **🗸** | **🗸** |  | 2 | 2 |
| **AVP24-10** |  | **🗸** | **🗸** | **🗸** | **🗸** |  | 4 | 2 |
| **AVP24-11** |  |  |  |  |  |  | 5 | 4 |
| **AVP24-12** |  |  |  | **🗸** | **🗸** |  | 3 | 6 |
| **AVP24-13** | **🗸** | **🗸** | **🗸** | **🗸** | **🗸** |  | 2 | 4 |
| **AVP24-14** |  |  | **🗸** |  |  |  | 0 | 4 |
| **AVP44-0** |  | **🗸** |  | **🗸** |  |  | 2 | 0 |
| **AVP44-1** | **🗸** | **🗸** | **🗸** |  |  |  | 1 | 0 |
| **AVP44-2** |  | **🗸** |  |  |  |  | 1 | 1 |
| **AVP44-3** |  |  | **🗸** | **🗸** |  | **🗸** | 5 | 4 |
| **AVP44-4** |  | **🗸** |  |  |  | **🗸** | 1 | 1 |
| **AVP44-5** |  |  | **🗸** | **🗸** | **🗸** |  | 5 | 1 |
| **AVP44-6** |  |  |  | **🗸** |  |  | 1 | 0 |
| **AVP44-7** |  | **🗸** |  |  |  |  | 0 | 0 |
| **AVP44-8** |  | **🗸** |  |  |  | **🗸** | 2 | 0 |
| **AVP44-9** | **🗸** | **🗸** | **🗸** |  |  | **🗸** | 0 | 2 |
| **AVP44-10** |  | **🗸** |  |  |  |  | 2 | 1 |
| **AVP63-0** |  |  |  |  | **n/a** | **n/a** | 9 | 3 |
| **AVP63-1** | **🗸** |  | **🗸** |  | **n/a** | **n/a** | 10 | 7 |
| **AVP63-2** |  |  |  |  | **n/a** | **n/a** | 10 | 5 |
| **AVP63-3** | **🗸** |  |  |  | **n/a** | **n/a** | 16 | 10 |
| **AVP63-4** | **🗸** |  |  |  | **n/a** | **n/a** | 6 | 5 |
| **AVP63-5** | **🗸** |  |  |  | **n/a** | **n/a** | 14 | 4 |
| **AVP63-6** |  |  |  |  | **n/a** | **n/a** | 14 | 6 |
| **AVP63-7** |  | **🗸** |  |  | **n/a** | **n/a** | 10 | 4 |
| **AVP63-8** |  |  |  |  | **n/a** | **n/a** | 6 | 8 |
| **AVP63-9** | **🗸** | **🗸** | **🗸** |  | **n/a** | **n/a** | 5 | 2 |
| **AVP63-10** | **🗸** | **🗸** |  |  | **n/a** | **n/a** | 7 | 5 |
| **AVP63-11** | **🗸** |  |  |  | **n/a** | **n/a** | 12 | 11 |
| **AVP63-12** |  |  |  |  | **n/a** | **n/a** | 11 | 1 |
| **AVP63-13** | **🗸** |  |  | **🗸** | **n/a** | **n/a** | 2 | 0 |
| **AVP63-14** | **🗸** | **🗸** |  |  | **n/a** | **n/a** | 3 | 10 |
| **AVP63-15** | **🗸** |  |  |  | **n/a** | **n/a** | 4 | 7 |
| **AVP63-16** | **🗸** |  | **🗸** |  | **n/a** | **n/a** | 9 | 0 |
| **AVP63-17** | **🗸** | **🗸** |  |  | **n/a** | **n/a** | 4 | 2 |
| **AVP63-18** | **🗸** |  |  | **🗸** | **n/a** | **n/a** | 9 | 1 |
| **AVP63-19** | **🗸** | **🗸** | **🗸** | **🗸** | **n/a** | **n/a** | 16 | 5 |
| **AVP63-20** | **🗸** | **🗸** |  |  | **n/a** | **n/a** | 16 | 7 |
| **AVP63-21** | **🗸** | **🗸** |  |  | **n/a** | **n/a** | 12 | 7 |
| **AVP63-22** | **🗸** | **🗸** |  | **🗸** | **n/a** | **n/a** | 4 | 7 |
| **AVP63-23** |  | **🗸** |  |  | **n/a** | **n/a** | 12 | 3 |

Caption

Table S1. Summary of rhythmic and non-rhythmic parameters by ROI and mouse. All ROIs analyzed for this study across 4 mice are arrayed in rows. Checkmarks indicate the detection of statistically significant circadian rhythmicity (Cosinor fit p<0.05) for each of our major measures, for 48h DD and LD datasets. The final 2 columns list the # of rhythmic coherence relationships each ROI had with others from the same mouse. Note that for AVP10, a DD recording was incomplete and not included in the dataset. Also, discrete, unambigious Ca^2+^ waves were less common in AVP63, so waves were not counted for rhythmicity for this mouse.
