## Supplementary figures and images for "Arginine-Vasopressin Expressing Neurons in the Murine Suprachiasmatic Nucleus Exhibit a Circadian Rhythm in Network Coherence *In Vivo*"

### Fig S1

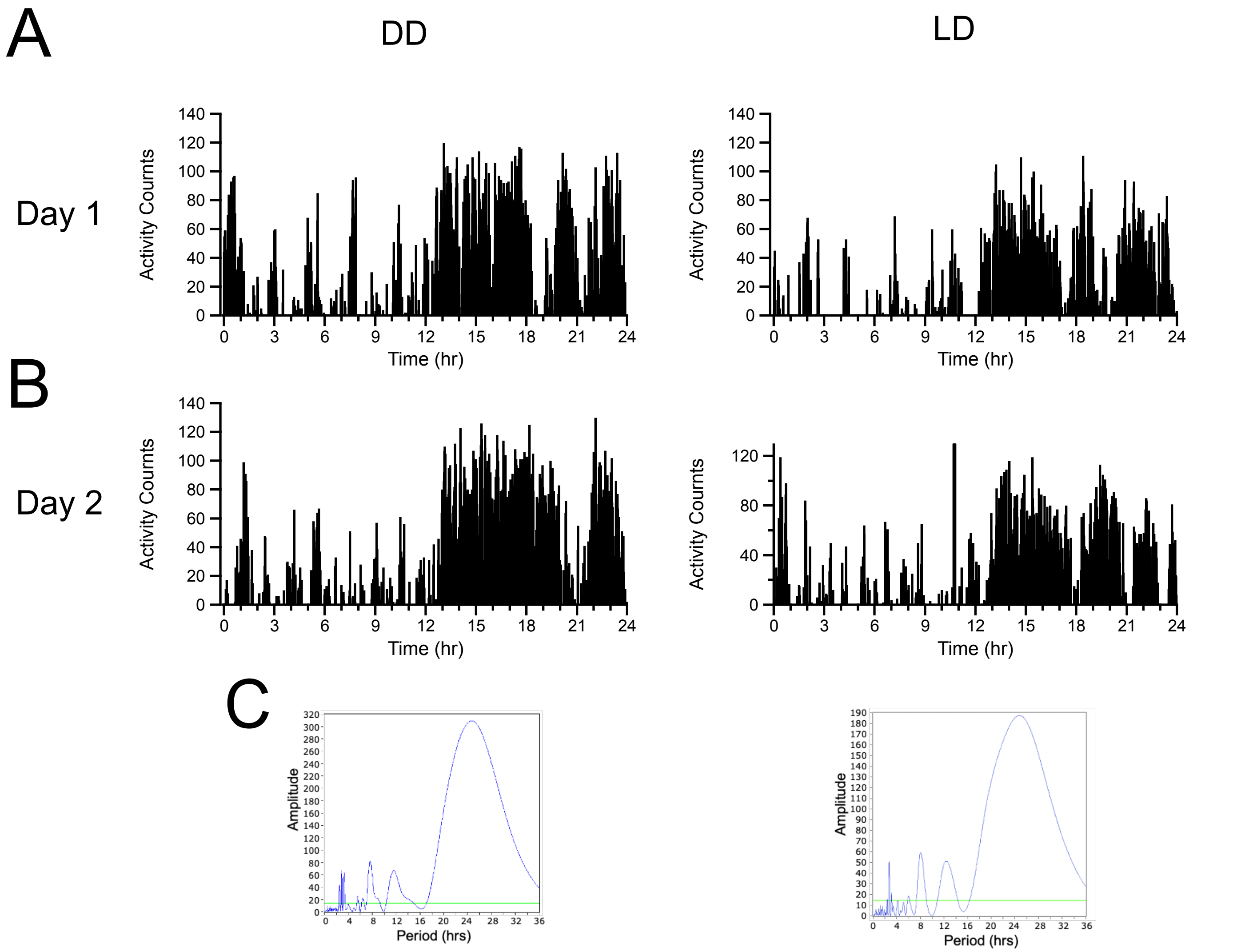

### Fig S2

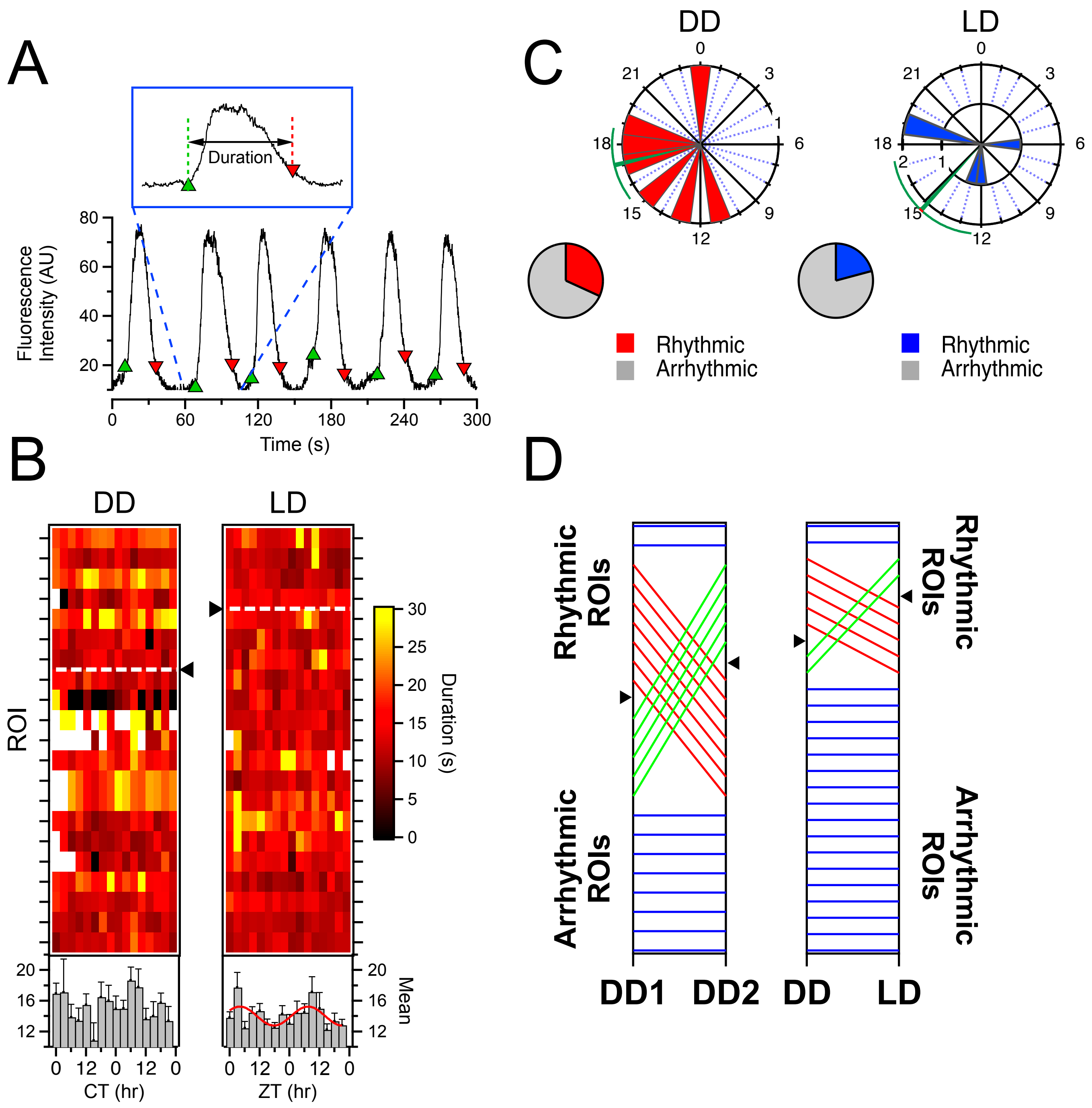

### Fig S3

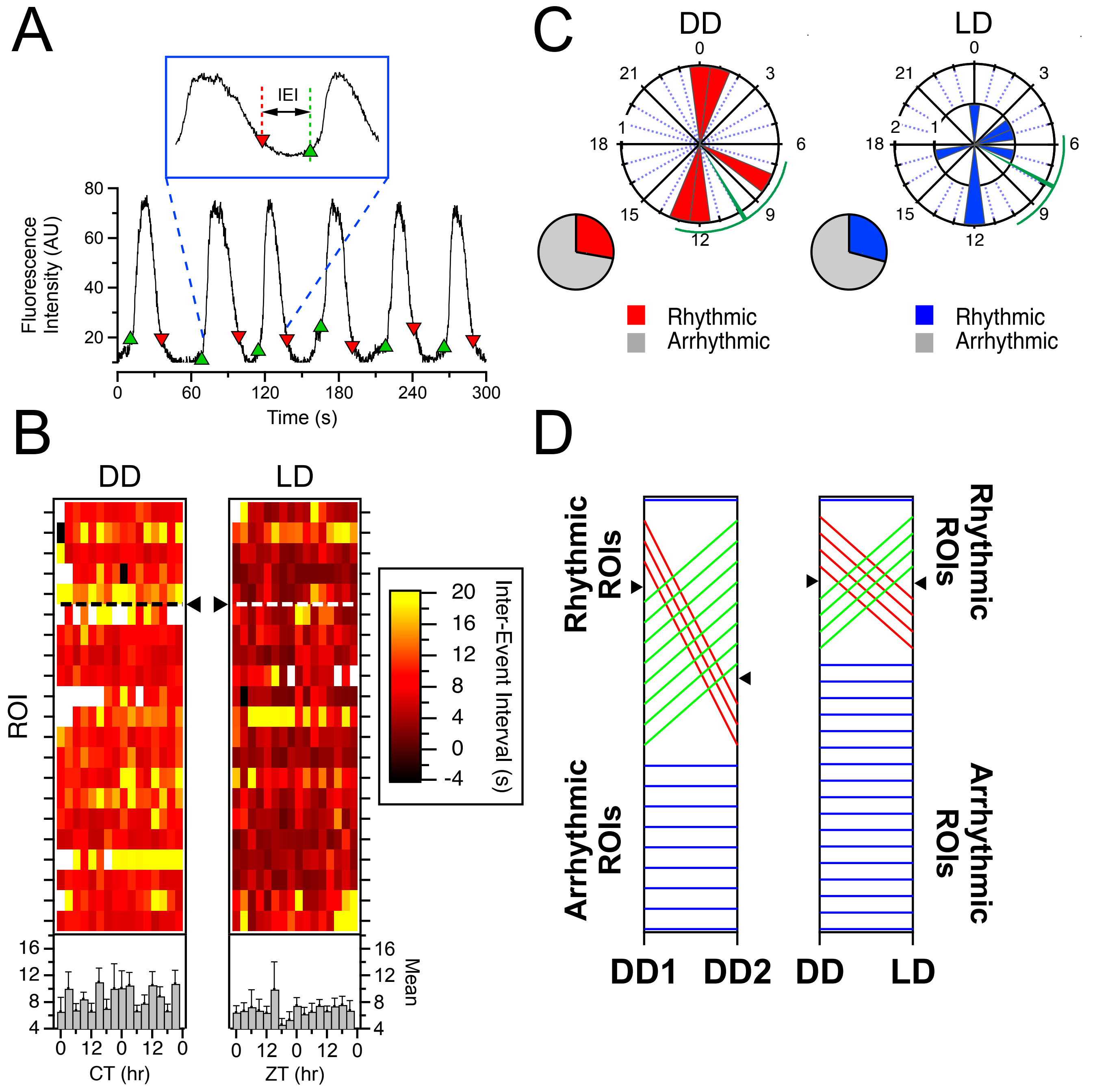

### Fig S4

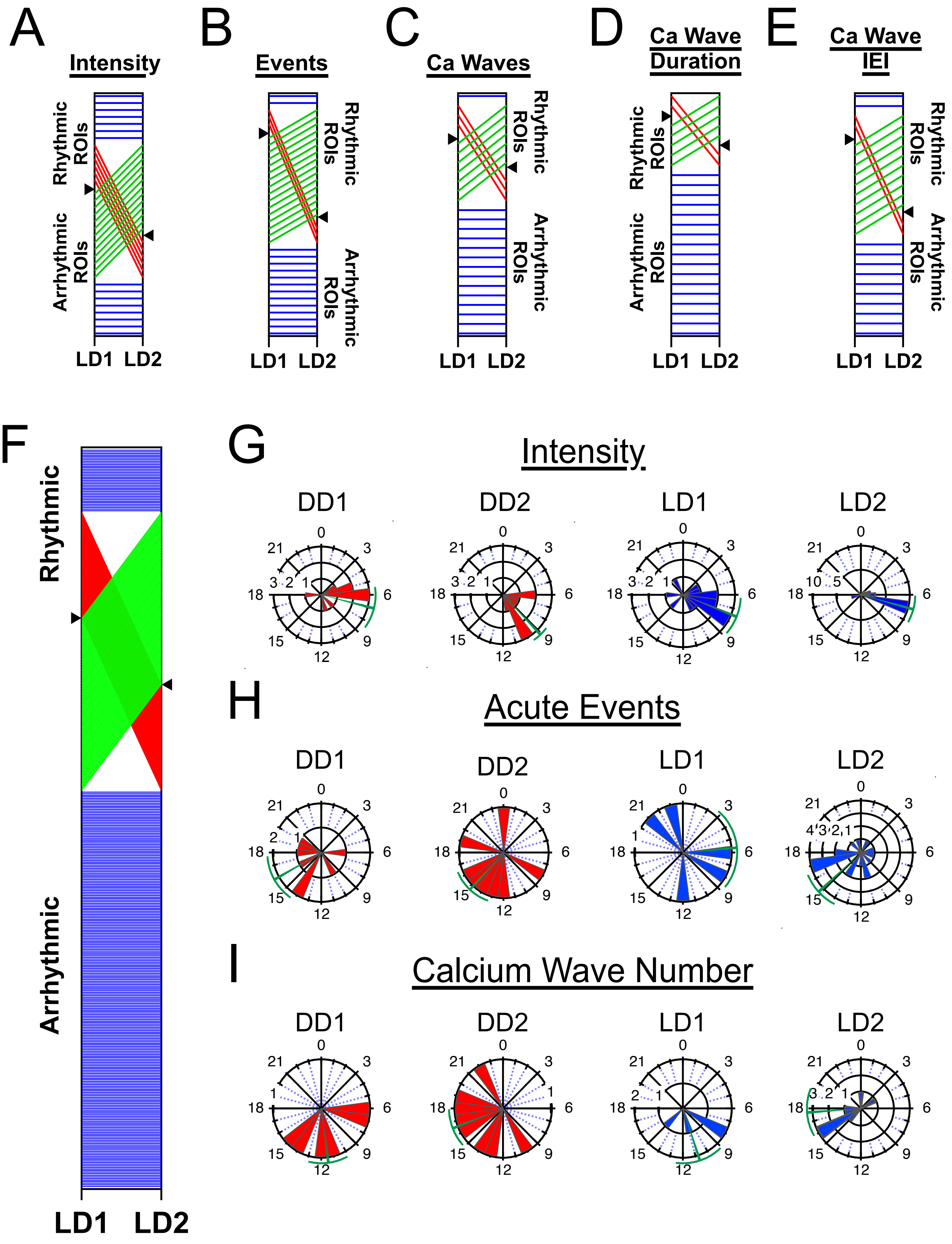

### Fig S5

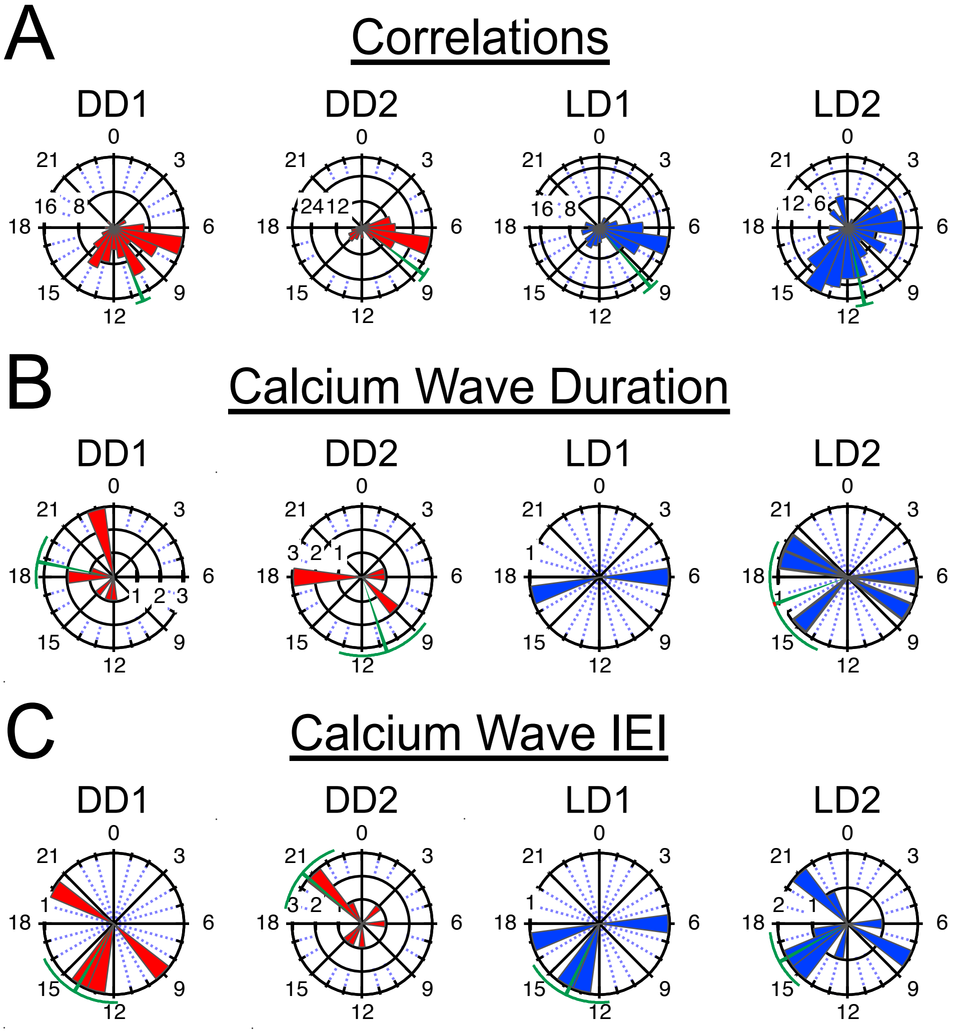

### Fig S6

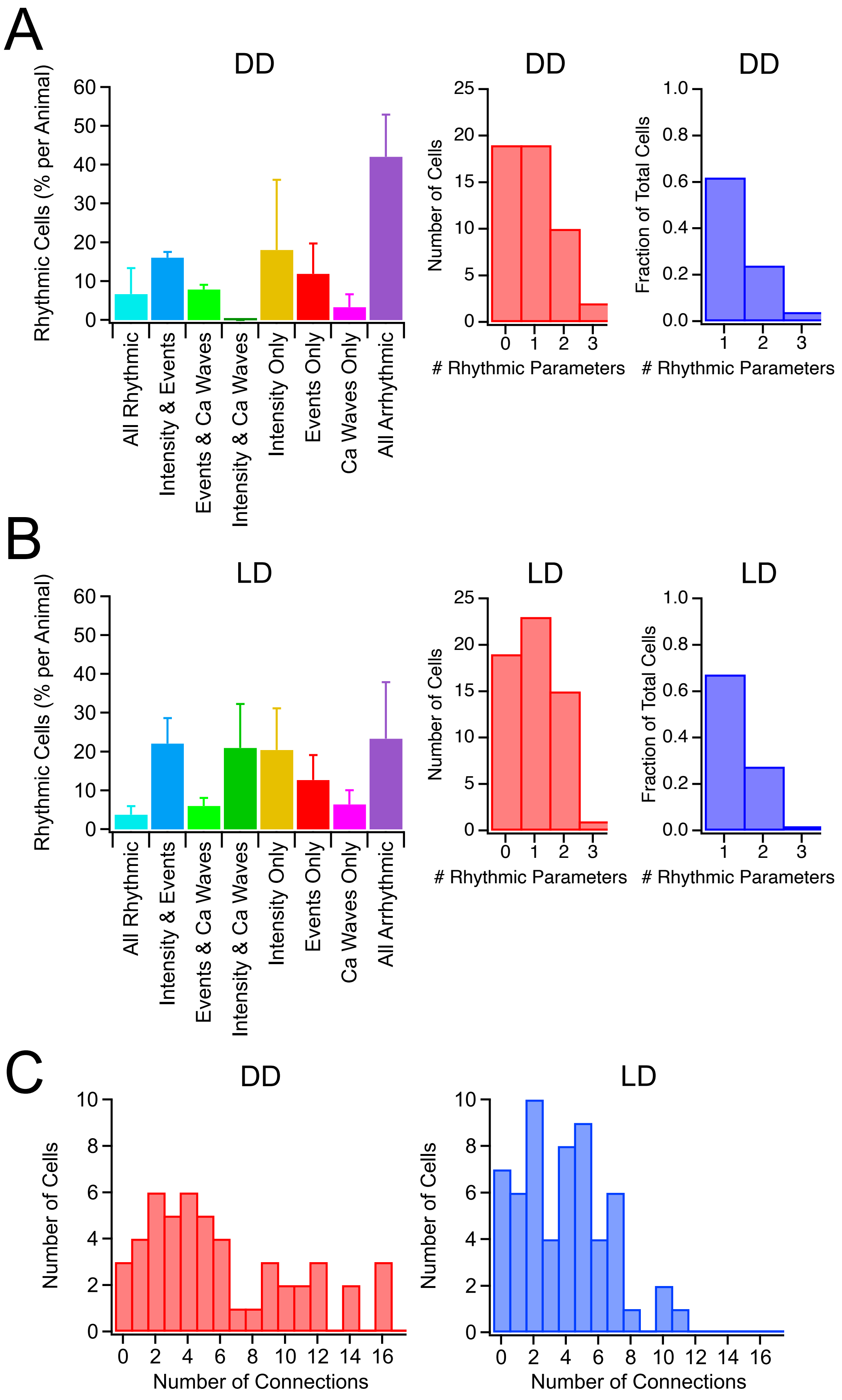
